# Reciprocal regulation of m^6^A modification and miRNA production machineries via phase separation-dependent and -independent mechanisms

**DOI:** 10.1101/2024.08.31.610644

**Authors:** Songxiao Zhong, Xindi Li, Changhao Li, Haiyan Bai, Jingjing Chen, Lu Gan, Jiyun Zhu, Taerin Oh, Xingxing Yan, Jiaying Zhu, Niankui Li, Hisashi Koiwa, Thomas Meek, Xu Peng, Bin Yu, Zhonghui Zhang, Xiuren Zhang

**Affiliations:** Department of Biochemistry and Biophysics, Texas A&M University, College Station, TX 77843, USA; Guangdong Provincial Key Laboratory of Biotechnology for Plant Development, School of Life Science, South China Normal University, Guangzhou 510631, China; School of Biological Sciences & Center for Plant Science Innovation, University of Nebraska-Lincoln, Lincoln, NE, USA; Biotechnology Research Institute, Chinese Academy of Agricultural Sciences, Beijing 10081, China; Department of Horticulture, Texas A&M University, College Station, TX 77843, USA; Department of Medical Physiology, College of Medicine, Texas A&M University, College Station, TX 77843, USA; Department of Biology, Texas A&M University, College Station, TX 77843

## Abstract

Methyltransferase complex (MTC) deposits *N*6-adenosine (m^6^A) onto RNA, whereas microprocessor produces miRNA. Whether and how these two distinct complexes cross-regulate each other has been poorly studied. Here we report that the MTC subunit B (MTB) tends to form insoluble condensates with poor activity, with its level monitored by 20S proteasome. Conversely, the microprocessor component SERRATE (SE) forms liquid-like condensates, which in turn promotes solubility and stability of MTB, leading to increased MTC activity. Consistently, the hypomorphic lines expressing SE variants, defective in MTC interaction or liquid-like phase behavior, exhibit reduced m^6^A level. Reciprocally, MTC can recruit microprocessor to *MIRNA* loci, prompting co-transcriptional cleavage of primary miRNA (pri-miRNAs) substrates. Additionally, pri-miRNAs carrying m^6^A modifications at their single-stranded basal regions are enriched by m^6^A readers, which retain microprocessor in the nucleoplasm for continuing processing. This reveals an unappreciated mechanism of phase separation in RNA modification and processing through MTC and microprocessor coordination.

## Introduction

Biomolecules such as intrinsically disordered proteins (IDPs) and nucleic acids can form discrete condensates and membrane-less compartments within a cellular or solute milieu^1^. IDPs typically contain unfolding regions that might serve as dock points for homotypic or heterotypic multivalent interactions, instigating phase separation or co-phase separation^2^. Liquid-liquid phase separation (LLPS) can amplify enzymatic reaction^3^, facilitate transcription^4,5^ and RNA metabolism^6,7^ among other biological processes^3,8^. Nonetheless, biomolecules may also assemble comparatively inert or inactive condensates. For example, IDPs can undergo spontaneous phase transition among heterogeneous behaviors, known as maturation and hardening, enabling functional switch^1,9^. To overcome these disorders or constraints, cells can deploy molecular chaperones for maintaining protein conformation and proteasomes for eliminating the misfolded proteins^10^. The plant kingdom encodes a multitude of IDPs, yet only a tiny fraction has been studied concerning their phase behaviors and physiological relevance^11,12^.

SE, the plant homolog of mammalian arsenic resistance protein 2 (ARS2), is a multi-functional IDP that acts in transcription, miRNA biogenesis, and pre-mRNA splicing among other processes^13^. SE also undergoes LLPS to facilitate the processing of pri-miRNAs by microprocessor and shuttling of miRNA/* duplexes to RNA-induced silencing complex^6,14^. Conversely, two other IDPs, SAID1/2, can form co-condensates with SE and sequester pri-miRNAs from microprocessor to inhibit miRNA production^15^. Notably, mis-folded or unsheltered SE is degraded by 20S proteasome^16,17^. These findings illuminate that SE, through its intrinsically disordered properties, serves as an assemblage nexus for multiple ribonucleoprotein (RNP) complexes to orchestrate above-mentioned RNA processing events^13^. This notwithstanding, it remains unclear whether and how SE/ARS2 contribute to additional RNA metabolism exemplified by epi-transcriptome modifications.

The mRNA adenosine methylase A (MTA) and MTB, plant homologs of mammalian methyltransferase like 3 (METTL3) and METTL14, respectively, function as essential constituents of MTC that catalyze m^6^A deposition. METTL3, once fused with a low-complexity domain of *Arabidopsis* photoreceptor protein CRY2, displays punctuated foci in mammalian cells^18^. In *Arabidopsis*, MTA seems to undergo co-LLPS with CRY2, granting light regulation of m^6^A of circadian clock associated transcripts^19^. Intriguingly, m^6^A itself can also serve as a scaffold facilitating the phase separation of RNP complexes^20,21^. In eukaryotes, m^6^A level is positively correlated with miRNA amount^22–24^. The m^6^A modification might locate in a stem-loop region, triggering a structural switch of pri-miRNAs that would be recognized by an RNA binding protein, and enabling the recruitment of DGCR8 for processing^24,25^. Alternatively, METTL3/MTA can recruit DGCR8 or TOUGH to facilitate pri-miRNA processing^22,23^. However, whether and how MTB undergoes the regulation of phase behaviors is unknown. In addition, the precise mechanism underlying MTC regulation of pri-miRNA processing remains unclear.

Here, we identified a non-canonical function of SE in promoting MTC-mediated m^6^A deposition. Unlike well-folded METTL14, MTB is an IDP and prone to forming insoluble condensates. However, SE could utilize its liquid-like phase behaviors to maintain the MTB solubility, allowing the uptake of RNA substrate by MTC to fulfill the enzymatic activity. Moreover, co-condensation of SE and MTB could also mutually protect each other from degradation by 20S proteasome. Reciprocally, MTC could tether the microprocessor to chromatin, facilitating co-transcriptional pri-miRNA processing. Furthermore, MTC could deposit m^6^A onto single-strand (ss) basal regions of pri-miRNAs, enabling m^6^A readers to retain microprocessor for further processing in the nucleoplasm. Thus, this study reveals how the two IDPs fine-tune their phase behaviors to regulate the epi-transcriptome modification, and how the two otherwise separate complexes orchestrate each other to promote miRNA production.

## Results

### SE displays spatiotemporal and physical association with MTC

To explore SE’s additional functions, we conducted a pan co-expression network analysis for principal components of microprocessor and MTC using over 1000 RNA-seq datasets^26^ (Extended Data Fig. 1a, Supplementary table 1). *SE*, *HYL1*, and *TOUGH* mutually exhibited moderate correlations, whereas their coordination with *DCL1* was less pronounced. Importantly, the substantial correlations emerged among *MTA*, *MTB*, and *SE*. Another MTC subunit, *FK506-BINDING PROTEIN 12 KD INTERACTING PROTEIN 37KD* (*FIP37)*, also showed a robust correlation with *SE*. Furthermore, *MTA* and *MTB* displayed correlation to other microprocessor components, to a certain extent, but not with the control gene, *Actin*. Additionally, *SE* and *MTC* depicted a co-expression pattern throughout various tissue (Extended Data Fig. 1b). To investigate if SE and MTC functioned genetically in a parallel or cooperative manner, we generated knockdown mutants for *MTA*, *MTB*, and *FIP37* (Extended Data Fig. 2a–f). All MTC mutants displayed developmental defects reminiscent of *se*, exemplified by upward-curly and narrow true leaves on one-week-old seedlings, as well as notched and trident-like, irregularly curled leaves, and stunted growth on three-week-old plants (Fig. 1a, Extended Data Fig. 2g). Subsequent RNA-seq profiling revealed a significant overlap of differential expressed genes (DEGs) between *se-2* and *mta* (Fig. 1b, Supplementary table 2). Moreover, yeast two hybrid (Y2H) screening assays showed that SE directly interacted with MTB and FIP37, but not with MTA (Fig. 1c, Extended Data Fig. 2h). Additionally, co-immunoprecipitation (co-IP), luciferase complementation imaging (LCI), and bimolecular fluorescence complementation (BiFC) (Fig. 1d, Extended Data Fig. 2i–m) further validated SE association with MTC. All these results suggested that SE and MTC might function within a shared genetic pathway.

**Figure 1.**
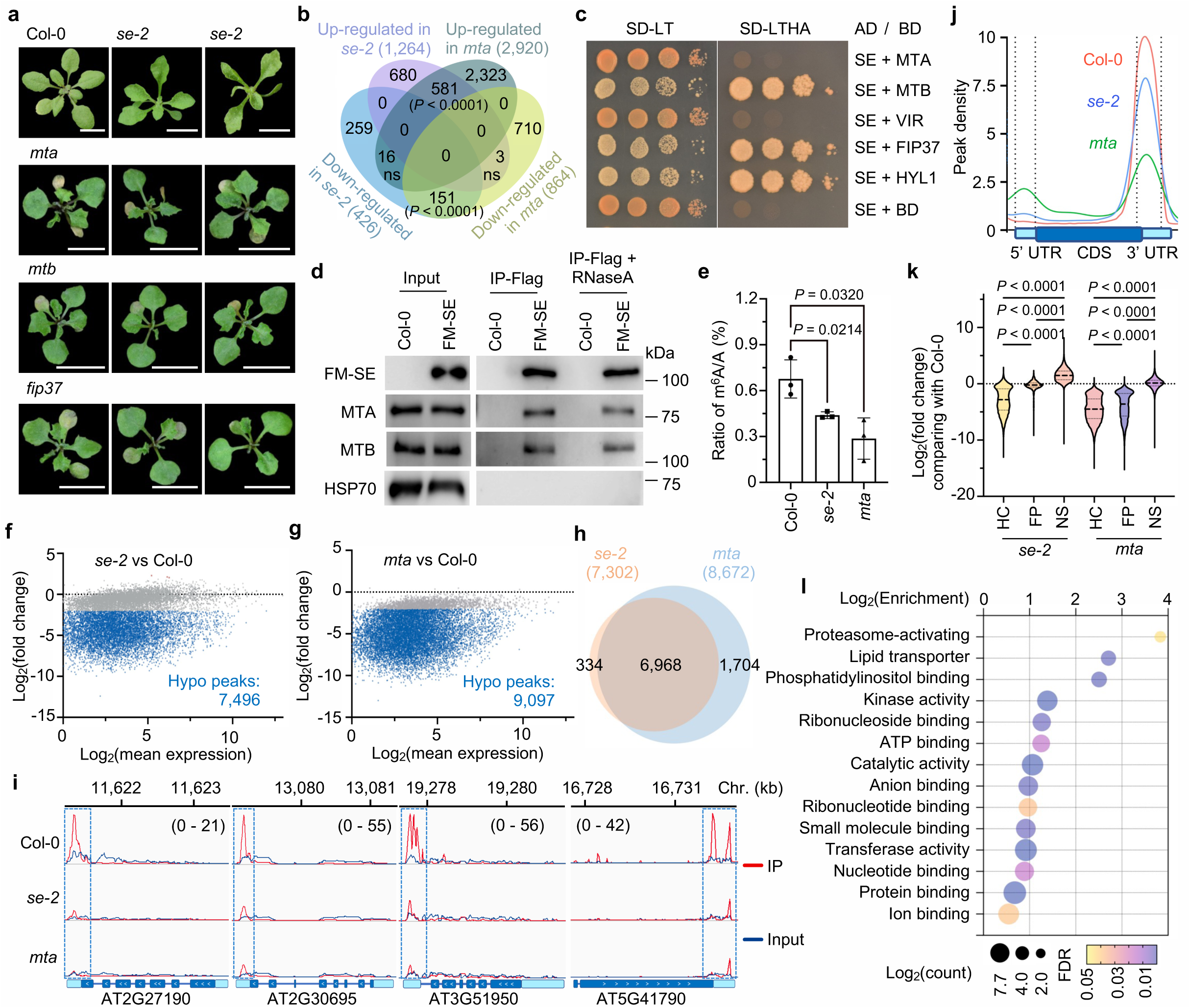
SE promotes m^6^A modification in *Arabidopsis*. **a,** *mta*, *mtb*, and *fip37* displayed developmental defects in three-week-old plants. Scale bar, 1 cm. **b,** Venn diagram illustrated significant overlap of DEGs between *se-2* and *mta* mutants vs Col-0. **c,** Y2H screening identified MTB and FIP37 as additional partners of SE. VIR, VIRILIZER. -LT, lacking Leu and Trp; -LTHA, lacking Leu, Trp, His and Ade. **d,** Co-IP assays validated the interaction between SE and MTC in the stable transgenic plants of Col-0; *pSE*::*Flag-4×MYC(FM)-SE*. IPs were conducted with anti-Flag antibodies. Immunoblots were detected anti-Flag or endogenous antibodies, respectively. HSP70 was a negative control. **e**, HPLC-MS analysis revealed a significant reduction of m^6^A global abundance in *se-2* and *mta* vs Col-0. Data are mean ± s.d. of three independent experiments. **f, g,** MA plots showed the differentially methylated peaks in *se-2* (**g**) and *mta* (**h**) vs Col-0. The red, blue and grey dots represent hypermethylated (log2(fold change) > 2, P < 0.01), hypomethylated (log_2_(fold change) < -2, P < 0.01), and comparable methylated m^6^A peaks, respectively. **h,** Venn diagram showed the overlap of hypomethylated genes from MeRIP-seq between *se-2* and *mta* when compared to Col-0. **i,** Integrative Genomics Viewer (IGV) files of selected hypomethylated genes. IP and input were represented in red and blue, respectively. Chr., chromosome. **j,** Metagene profile showed m^6^A distribution cross the gene body, with x and y-axes representing relative position and abundance of m^6^A modification on transcripts, respectively. **k,** Violin plot showed peak differential methylation levels enriched in mutants relative to Col-0. The lines represent the median and quartiles. HC, High-confidential-m^6^A peaks; FP, false-positives-m^6^A peaks; NS, non-significant-m^6^A peaks. **l,** GO analysis of shared hypomethylated genes in mutants vs Col-0. At least fifteen independent transgenic lines were photographed (**a**), ten independent colonies (**c**), three biological replicates (**d**) were tested, and representative images are shown. Hypergeometric distribution test (**b**, **l**), unpaired two-sided t-test (**e**), DESeq2 (**f**, **g**), and one-way ANOVA with Dunnett’s multiple comparison test (**k**). P values are shown in panels (**b**, **e**, and **k**).

### SE impacts transcriptome-wide m^6^A deposition

We next assessed the impact of SE on m^6^A deposition through high-performance liquid chromatography tandem mass spectrometry (HPLC-MS). We quantified m^6^A amount of poly(A)+ RNAs with two controls (Extended Data Fig. 3a). Indeed, *se-2*, like *mta*, displayed a significant reduction of m^6^A level vs Col-0 (Fig. 1e). Next, we conducted methylated RNA immunoprecipitation (MeRIP) assays (Extended Data Fig. 3b). Whereas the starting amount of RNA input was comparable cross the samples, MeRIP-enriched signals of m^6^A-harbored fragments were clearly reduced for *se-2* and *mta* mutants vs Col-0 (Extended Data Fig. 3c), indicative of reduced levels of methylated mRNAs from the mutants vs Col-0.

To quantitatively compare the m^6^A epi-transcriptome across different samples, we recalled over 50,000 m^6^A candidate peaks via exomePeak2 software with a setting of log_2_(enrichment) > 0 (Supplementary table 3). We defined high-confidence (HC)-m^6^A peaks of interest using a threshold of P < 0.05 and log_2_(enrichment) > 1, while the rest of peaks were claimed as non-significant (NS)-m^6^A peaks and used as controls for comparative analysis with DEseq2. In this scenario, we identified 11,487 HC-m^6^A peaks in Col-0, wherein a substantial proportion of the peaks were hypomethylated in *se-2* and *mta* (Fig. 1f,g, Supplementary table 3). Specifically, 7,496 and 9,097 peaks were hypomethylated on 7,302 and 8,672 transcripts in *se-2* and *mta*, respectively, with 6,968 in common (Fig. 1h,i). Meta-analysis of m^6^A-enriched reads showed reduced distributions at 3’ end of transcripts (Fig. 1j). A motif analysis of the hypo-peaks revealed the presence of canonical m^6^A motif, RRACH context (R = A and G; H = A, C, and U) (Extended Data Fig. 3d), and the plant-specific URUAY (Y = C and U) motif^27^.

Similarly, we obtained a total of 11,963 and 11,667 seemingly “HC”-m^6^A peaks in *se-2* and *mta*, respectively, with 1,494 and 3,309 peaks enriched specifically in the according mutants (Supplementary table 3). These peaks are likely false positives (hereafter referred to FP-m^6^A peaks) in part due to the internal normalization algorithm within the samples, given that the overall m^6^A level on FP-m^6^A peaks was reduced in the mutants vs Col-0 (Fig. 1k). Furthermore, Go-Ontology (GO) analysis showed that the m^6^A transcripts co-regulated by SE and MTA were significantly enriched in metabolic processes, proteasome-activating, and lipid transporter pathways (Fig. 1l). Taken together, these results indicated that SE may promote MTC-mediated m^6^A deposition.

### IDRs confer the SE-MTB interaction

We employed an *in-silico* approach to explore the structural basis underlying the MTB-SÈs contribution to m^6^A deposition. Both IUPred3^28^ and neural network-based^29^ bioinformatic analyses suggested that plant MTBs, unlike their counterparts in metazoans and fungi, had largely extended N-terminal intrinsically disordered regions (IDRs, up to 600 amino acids), whereas plant MTAs exhibited a comparable feature to mammalian METTL3 (Fig.2a–c Extended Data Fig. 4a–e). A three-dimensional (3D) simulation of the MTA-MTB heterodimer via AlphaFold2-multimer^30^ showed that the core domains assembled in an architecture reminiscent of human MTC^31^ (Extended Data Fig. 4g). Subsequently, the simulation of SE-MTB assembly revealed that IDRs of two proteins intermingled each other (Extended Data Fig. 4h) with a majority of interactive hydrogen bonds harbored in their IDRs (Extended Data Fig. 4b–e). Indeed, Y2H assays validated that the C-terminal region (469 to 720 aa) of SE interacted with the N-terminal IDR (1 to 634 aa) of MTB (Fig. 2f). Computation analysis predicted that seven putative amino acids within the C-terminal region of SE may serve as donors for interactive hydrogen bonds with MTB (Extended Data Fig. 4i). Y2H assays with alanine scanning variants of seven potential hydrogen bond donors (562, 580, 584, 586, 589, 666, and 718) pinpointed R718 as a critical residue for SE’s interaction with MTB (Fig 2f,g), but not with DCL1. Importantly, R718 was depleted in all *se* mutants (Fig. 2h)

**Figure 2.**
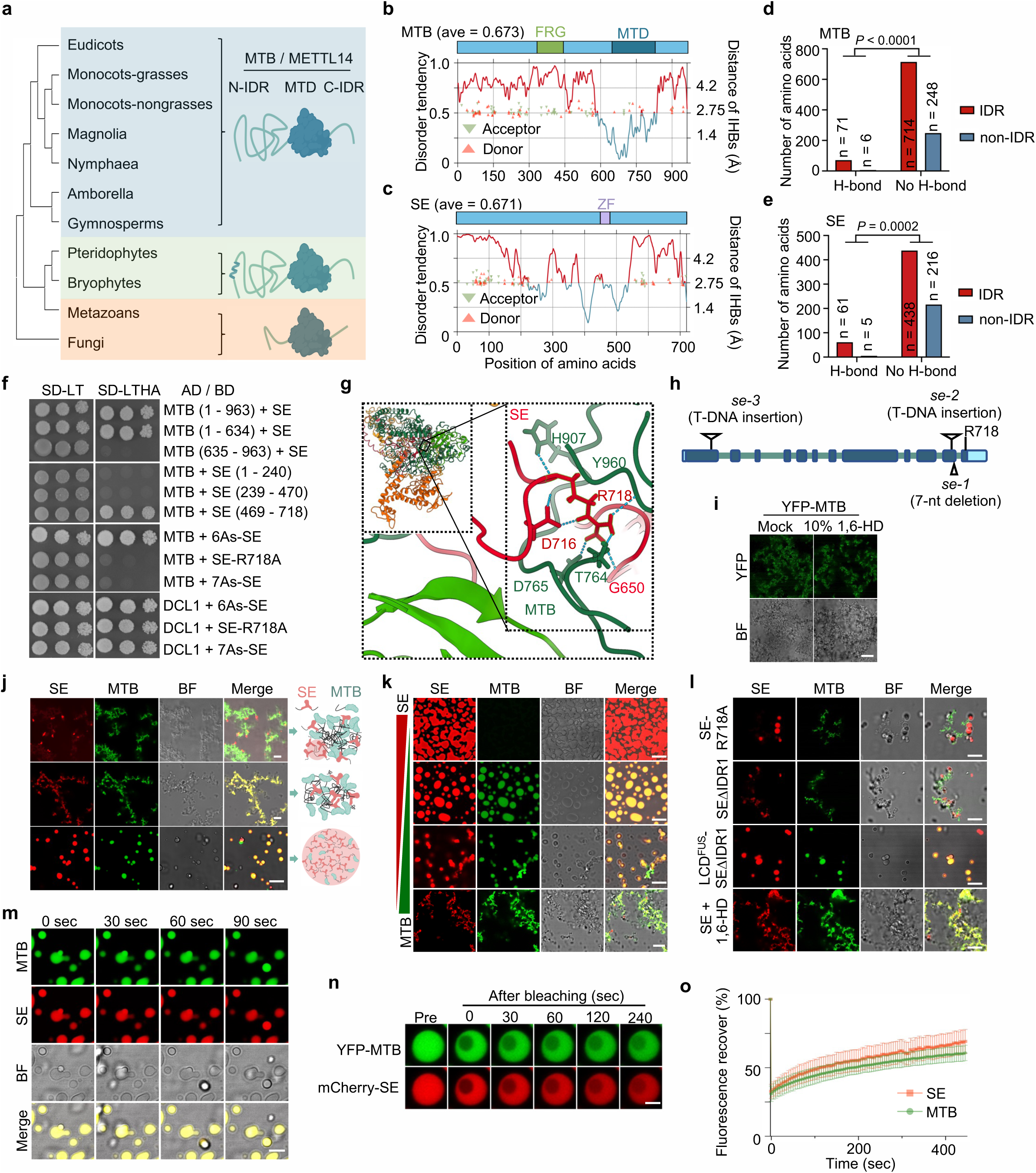
SE modulates the phase behaviors of MTC *in vitro*. **a,** Phylogenetic tree showed that plant MTBs have extended N-terminal IDRs. **b**,**c,** Computational simulation predicted disordered (red) and ordered (blue) regions of MTB (**b**) and SE (**c**) and their interactive hydrogen bonds (IHBs). Triangles and right y-axis, IHBs and their corresponding distances, respectively. MTD, methyltransferase domain. ZF, zinc finger; FRG, (F/Y)RG motif enriched region. **d, e,** Simulation indicated the significant enrichment of IHBs in the IDRs of MTB (**d**) and SE (**e**) in the complex. P values, two-sided fisher’s exact test. **f,** Fine-mapping of SE-MTB interaction regions. 7As-SE, alanine substitutes of all seven potential IHB donors in SE; 6As-SE, a variant resembled 7As-SE except R718. Refer to Fig.1c for legend. **g,** Simulation identifies R718 as key in SE-MTB interaction. **h,** Illustration of alleles of *se* mutants. **i**, *In vitro* phase behaviors of 3 μM YFP-MTB in the droplet buffer with or without 10% 1,6-HD. **j,** Confocal microscopy detected three non-exclusive SE-MTB co-condensate scenarios. The top, middle and bottom panels showed MTB aggregates encapsulated inside a SE shell, the co-aggregation, and fully overlapping spherical condensates, respectively. 1 μM of SE and 0.5 μM (top and middle) or 0.1 μM (bottom) of MTB were used. **k,** Confocal microscopy of SE-MTB *in vitro* co-condensation at different ratios. SE vs MTB (μM), from top to bottom, 10 vs 0, 5 vs 0.15, 1.5 vs 0.5, and 0.5 vs 3. **l,** Confocal microscopy shows the dependance of SE-MTB co-condensation on SE’s IDR1 and its interaction with MTB *in vitro*. SE-R718A, which has compromised interaction with MTB; SEΔIDR1, which lacks the N-terminal IDR; LCD^FUS^-SEΔIDR1, a chimeric protein of human FUS’s LCD and SEΔIDR1. **m,** Time-lapse live images showed the fusion of SE-MTB co-condensates. **n,o,** FRAP analysis showed the mobility of SE-MTB condensates *in vitro*. Data are mean ± s.d. of eight independent experiments. At least ten colonies (**f**), three (**i** – **m**) and eight (**o**) independent experiments were performed, and representative images are shown. Scale bars, 10 μm (**i**, **m**), 5 μm (**j**, **k**, and **l**), and 2.5 μm (**n**).

### SE modulates phase behaviors of MTB *in vitro*

Since SE and MTB are IDPs and interact with each other, we next investigated their phase behaviors. Recombinant SE indeed formed spherical condensates (Extended Data Fig. 5a,b). This phase behavior relied on the N-terminal IDR (IDR1) as its depletion (SEΔIDR1) disrupted the condensation, and the defect could be rescued by the fusion of a low-complexity domain from mammalian FUS (LCD^FUS^-SEΔIDR1), as observed earlier^6^ (Extended Data Fig. 5a,b). Moreover, SE-R718A showed analogous LLPS capability to wild-type (WT) SE and LCD^FUS^-SEΔIDR1 (Extended Data Fig. 5a,b). This result indicated that the mutation of R718A did not alter SE phase behavior despite of compromising its interaction with MTB *in vitro*.

Recombinant MTB was soluble in a 500 mM salt buffer but turned turbid upon lowering the salt concentration to 150 mM (Extended Data Fig. 5c). Subsequent microscopic examination revealed irregular morphology of MTB condensates (Fig.2i). MTA remained soluble under low salt conditions but became susceptible to irregular condensate upon stimulation by a crowder (5% Ficoll) (Extended Data Fig. 5e). Notably, these condensates could not be disrupted with the addition of 10% 1,6-hexanediol (1,6-HD), indicative of insoluble states (Fig.2i, Extended Data Fig. 5d). In addition, the phase behaviors of neither MTA nor MTB were affected by fluorescence tag (Extended Data Fig. 5e).

The strong tendency of SE-MTB to form heterotypic assembly raised a question of whether SE could impact the phase behaviors of MTB. To test this, we conducted *in vitro* phase separation assays in different ways. We first reduced the salt concentration of the protein solutions to 150 mM NaCl, and then mixed the two proteins in the identical condition. Upon dilution, MTB promptly aggregated, leaving SE onto the surface of insoluble condensates, thereby co-condensation in irregular shape (Fig. 2j, top). Consequently, a few puncta comprising solely SE were observed (Fig. 2j, top). We next mixed the solution of two proteins prior to the buffer conversion. Similarly, such procedure also resulted in concurrent irregular condensates (Fig. 2j, middle). These observations prompted us to hypothesize if the initial formation of SE droplets is necessary to avert protein aggregation. To test this, a gradient amount of SE was used for droplet assembly before the addition of MTB (Fig. 2j, bottom). This time, upon mixing, SE and MTB simultaneously exhibited a spherical morphology when SE was 10-fold or more the amount of MTB (Fig. 2j, bottom). However, increasing the ratio of MTB/SE would readily cause the formation of irregular condensates (Fig. 2k). Moreover, the use of LCD^FUS^-SEΔIDR1, but neither SE-R718A nor SEΔIDR1, displayed spherical co-condensates (Fig. 2l). In addition, the human LCD^FUS^ condensates were unable to confer the liquid-like behavior to MTB (Extended Data Fig. 5f). These results indicated that SE droplets could modulate the phase behaviors of MTB, a process entailing direct interaction at the C-terminal of SE and the intrinsic LLPS properties conferred by the N-terminal IDR of SE. In contrast, mixture of MTA and MTB in this scenario formed exclusively irregular co-condensates (Extended Data Fig. 5g).

We further assessed the phase behavior of SE-MTB co-condensates. The 3D rendering images revealed that both SE and MTB indeed displayed a spherical shape (Extended Data Fig. 5h). The condensates were disrupted by addition of 1,6-HD (Fig. 2l, bottom). Importantly, the approaching droplets that consisted of two proteins were able to fuse into one larger condensate (Fig. 2m). In addition, FRAP assays showed the recovery of fluorescence after bleaching, indicative of the *bona fide* mobility of SE-MTB condensates (Fig. 2n,o). Taken together, the observations indicated that pre-existing SE liquid droplet would be a perquisite of controlling the phase behavior of MTB to prevent the protein aggregation *in vitro*.

### MTC displays SE-dependent LLPS *in vivo*

We next asked if SE could influence the phase behaviors of MTC *in vivo*. We co-expressed MTA-CFP, YFP-MTB, and mCherry-SE in mesophyll cells obtained from Col-0. The rendered 3D fluorescence revealed that the three proteins co-localized within discrete, spherical condensates (Extended Data Fig. 5i,j). Time-lapse live imaging demonstrated that two approaching condensates could fuse into one larger body (Extended Data Fig. 5k). Furthermore, FRAP assays revealed that both signals regained within a two-minute timeframe, suggesting a dynamic fluidity intrinsic to the SE-MTC co-condensates *in vivo* (Extended Data Fig. 5l,m).

In parallel, we prepared protoplasts from *se-1* in which SE protein is negligibly presented. In this situation, MTA-CFP and YFP-MTB also exhibited condensates (Extended Data Fig. 5n). However, these condensates could not recover after bleaching in FRAP assays (Fig. 3a), indicative of gel-like or solid-like phase behaviors, reminiscent of their *in vitro* patterns (Extended Data Fig. 5f) and also some other RNPs *in vivo*^32–34^. Again, the supplement of WT SE and LCD^FUS^-SEΔIDR1, but not SE-R718A and SEΔIDR1, could confer fluid properties to the co-condensates as the fluorescence of the former, but not the latter condensates, was quickly recovered after the photobleaching as observed in Col-0 (Fig. 3a, Extended Data Fig. 5n). Additionally, treatment of 1,6-HD disrupted the SE-granted mobility of MTC condensates, reflecting by hardening of the complex (Fig. 3a, Extended Data Fig. 5n).

**Figure 3.**
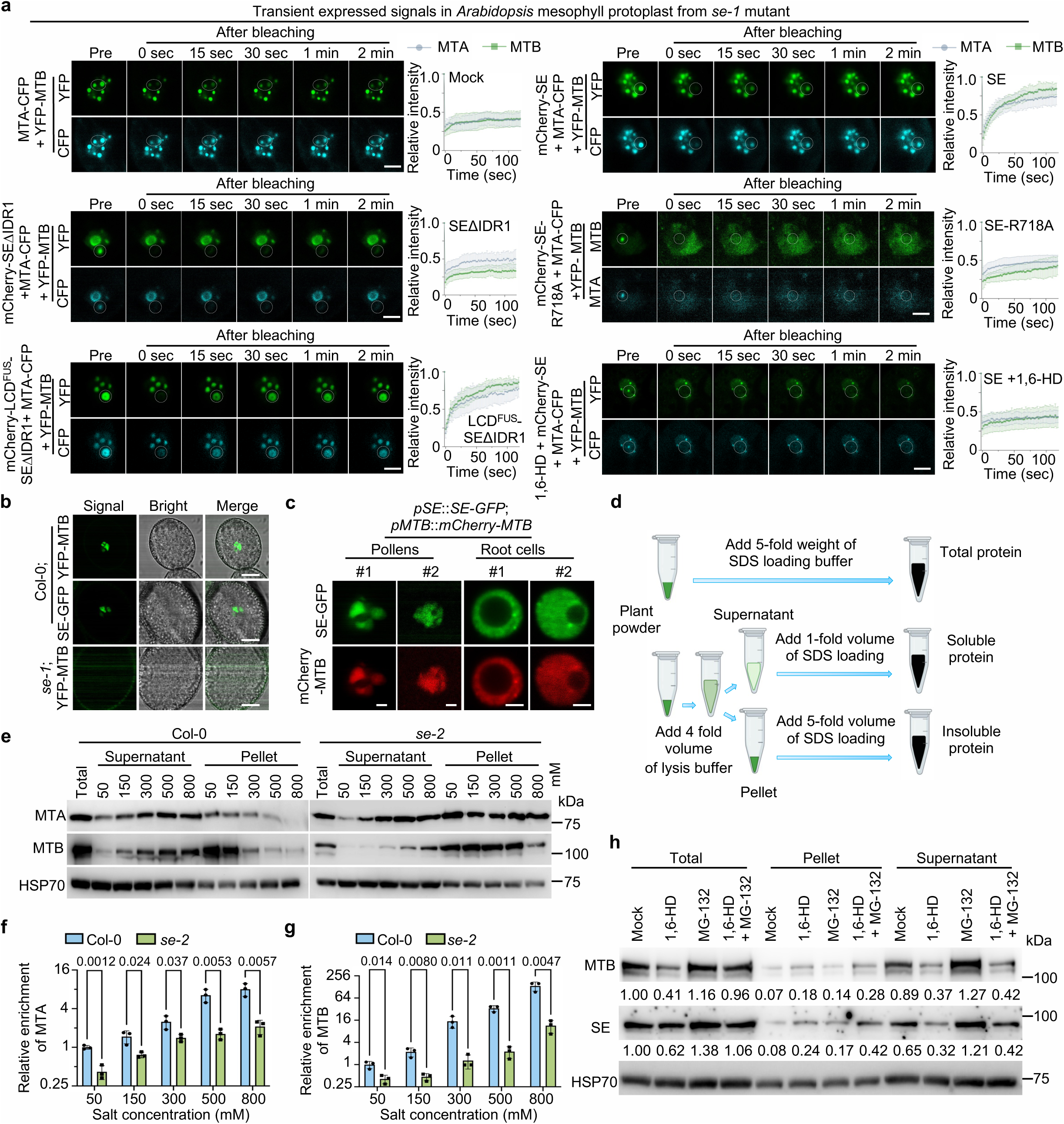
SE confers the liquid-like phase behaviors to MTC *in vivo*. **a,** FRAP assays showed that transiently expressed proteins in mesophyll cells from *se-1*, supplemented with or without different forms of SE variants, display distinct phase behaviors. SE-R718A, SEΔIDR1, and LCD^FUS^-SEΔIDR1 refer to Fig.2l. **b, c,** Confocal microscopy images showed that SE-dependent MTB condensates in pollens of transgenic plants (**b**) and the SE and MTB co-condensates in both somatic cells and pollens (**c**). **d,** Schematic illustration of the protein extraction from plant powder for sedimentation assay. See method section for detailed description. **e – g,** Immunoblots (**e**) and statistical analysis of sedimentation assays demonstrated the obvious enrichments of both MTA (**f**) and MTB (**g**) proteins in the insoluble fraction from *se-2* vs Col-0. HSP70 was a loading control. The ratio was calculated as (soluble MTB / soluble HSP70) / (insoluble MTB / insoluble HSP70) and was normalized to the Col-0 value at 50 mM salt concentration, which is set to 1. Unpaired two-sided t-test. The numbers above columns in (**f**, **g**) are indicated p values. **h,** Immunoblots detected the protein solubility upon chemical treatments. Ten-day-old seedlings were used. Note: both 1,6-HD and co-supply of 1,6-HD and MG-132 triggered the transition of SE and MTB complex from a soluble form to aggregates, compared with mock and MG-132 treatment, respectively. Data are mean ± s.d. of eight (**a**) and three (**f**, **g**) independent experiments. At least eight independent protoplasts and cells (**a**, **b**, and **c**), three independent experiments were performed (**e**, **h**), and representative images are shown. Scale bars, 5 μm (**a**, **b**), and 2.5 μm (**c**).

Furthermore, we transiently expressed MTC and SE under 35S promoter or their native promoters in *N. benthamiana*. In either case, SE and MTC could form co-condensates, despite the lower expression under the native promoters led to fewer and smaller co-condensates than that of 35S promoter (Extended Data Fig. 6a). Notably, the co-condensates exhibited similar mobility regardless of protein levels *in vivo* (Extended Data Fig. 6b–d). Finally, we assessed phase separation of native MTB that was expressed via its own promoter in stable transgenic plants. Again, MTB and SE could form co-condensates in both somatic cells and pollens (Fig. 3b,c). Clearly, the condensates displayed mobility as the fluorescence were recuperated within a 30-second duration in FRAP assays (Extended Data Fig. 6e). Furthermore, the application of 1,6-HD induced a phase transition to hardened condensates (Extended Data Fig. 6e). Importantly, the MTB signal intensity became weaker and the condensates disappeared when crossed into *se-1* (Fig. 2b, Extended Data Fig. 6f,g), indicative of SE-driven formation of MTC droplets in plants. Hence, SE was able to confer liquid-like properties to MTC *in vivo*, too.

### MTC exhibits SE-dependent soluble condensates in plants

To further study the impact of SE on the physical status of MTC, we evaluated the solubility of MTC in Col-0 and *se-2* plants by sedimentation analysis. In the assays, soluble proteins were prepared using lysis buffers containing a potent detergent and varying salt concentrations, whereas the resulting pellet after centrifuge, reflecting the insoluble protein fraction, was resuspended with SDS-loading buffer (Fig. 3d). Western blot analysis revealed that MTB, along with MTA but to a less extent, was significantly enriched in the insoluble fraction of *se-2* vs Col-0 (Fig. 3e–g). Meanwhile, the levels of both soluble MTA and MTB were reduced more significantly in the null-allele *se-3* (Extended Data Fig. 6h). Slightly different from the scenario in the adult plants (Extended Fig. 2c), *MTB* mRNA level, and resultant protein amount, slightly increased in young seedling of *se-2* vs Col-0 (Extended Fig. 6i). Despite this, the ratio of the soluble fraction vs the one in insoluble pellet remained decreased in the *se-2* vs Col-0 (Extended Data Fig. 6j). To further clarify if liquid-like properties were required for their solubility, we conducted 1,6-HD treatment prior protein extraction and found that both MTB and SE condensed into the pellets. We noticed that total proteins of SE and MTB were reduced after 1,6-HD treatment, and this reduction could be restored by co-supply of MG-132, in which the soluble protein contents were still decreased, while it became enriched in the pellet fraction (Fig. 3h). Collectively, these results suggested that SE-driven liquid-like phase behaviors of MTC are critical for the protein solubility in plants.

### SE facilitates m^6^A deposition

Since LLPS could mediate the exchange of molecules with the surrounding milieu, we hypothesized that SE modulation of MTA/MTB condensates might impact the incorporation of RNA in the complex. To test this, we introduced Cy5-labeled RNA into a mixture of SE droplets and protein solutions of MTC. We observed that the fluorescence signal of condensates, encompassing a tripartite protein ensemble in a spherical shape along with the RNA, exhibited substantial overlap (Fig. 4a). In contrast, when we treated the protein mixture with 1,6-HD before supply of RNA, signals displayed only partially overlapped (Fig. 4a), implying that RNA was lingered on the surface of irregular-shaped condensates *in vitro*. Meanwhile, we immunoprecipitated MTA from nuclear fractions and found that the enrichment of MTC-bound RNAs was reduced in *se-1* vs Col-0 (Fig. 4b). These results indicated that SE promotes RNA incorporation into MTC condensates. The prediction from this model would be that the compromised loading of RNA into MTC caused by the loss of SE might inhibit the MTC enzymatic activity. To test this, we conducted *in vitro* methylation assays with or without SE or its variants. After the reaction, we removed proteins and enriched methylated RNAs using an anti-m^6^A antibody, followed by P^32^ labeling and autographic imaging. Indeed, SE and LCD^FUS^-SEΔIDR1 caused significant increase in MTC activity, whereas SE-R718A and SEΔIDR1 had minimal effect, comparing to controls (Fig. 4c,d). Altogether, we concluded that SE could modulate the phase behaviors of MTC to facilitate m^6^A deposition.

**Figure 4.**
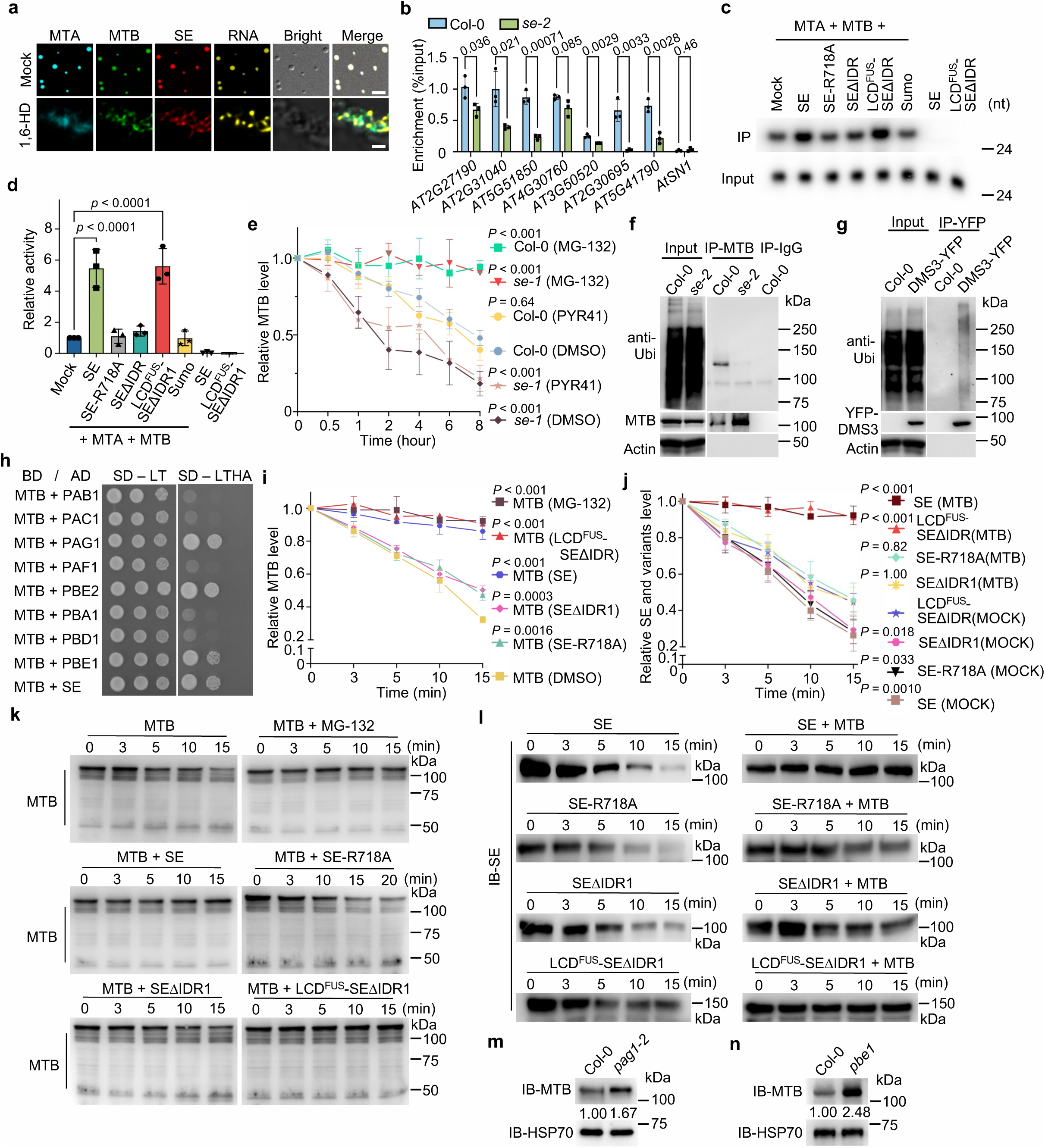
SE-dependent phase behavior of MTC is critical for its enzymatic activity and protein stability. **a,** Confocal microscopy examined the incorporation of proteins and RNA into condensates *in vitro* without (top) or with (bottom) 10% 1,6-HD. Scale bars, 5 μm (top) and 1 μm (bottom). **b,** MTA RIP-qPCR showed the decreased association between MTC and RNA in *se-2* vs Col-0. IPs were conducted with nuclear fraction using an anti-MTA antibody. The numbers above columns are indicated p values. **c, d,** *In vitro* m^6^A methylation assays (**c**) and the statistical analysis (**d**). **e**, MTB showed a shorter half-life in *se-1* vs Col-0 and the decay rate was hindered by MG-132, but not by PYR-41 or DMSO. **f, g,** Immunoblot assays of indicated immunoprecipitates did not detect poly(Ubi)-conjugated forms of MTB in planta. IPs were conducted using anti-MTB conjugated (**f**) or GFP-trap beads (**g**), respectively. IgG-IP, a negative control. Immunoblots were performed with indicated antibodies. **h,** Y2H assays showed the interactions between MTB and 20S proteasome subunits. Refer to Fig.1c for legend. **i – l**, 20S proteasome reconstitution decay assays showed that preincubation of MTB with mCherry tagged SE and LCD^FUS^-SEΔIDR1, but not SEΔIDR1 or SE-R718A, could mutually protect each other from degradation. The notations in and outside the brackets represent detected and co-incubated proteins, respectively. **m, n,** Immunoblots showed that MTB accumulates in *pag1-2* (**m**) and *pbe1* (**n**) vs Col-0 where the amount was arbitrarily assigned a value of 1. MTB was detected with endogenous antibodies. HSP70 was an internal control. For **c**, **d**, **i – l,** SE-R718A, SEΔIDR1, and LCD^FUS^-SEΔIDR1 refer to Fig. 2l. Data are mean ± s.d. of three independent experiments (**b**, **e**, **i**, and **j**). At least three independent experiments were performed (**a**, **c**, **f**, **g**, **k**, **l**, **m**, and **n**), ten colonies were detected (**h**), and representative images are shown. P values, unpaired two-sided t-test (**b**) and one-way ANOVA analysis with Tukey’s multiple comparisons test (**d**, **e**, **i**, and **j**). Only the comparisons with significance (**d**), the Col-0 (DMSO) (**e**), MTB (DMSO) (**i**), and LCD^FUS^-SEΔIDR1 (Mock) (**j**) groups are shown. For detailed analysis, see supplementary table 4.

### Co-condensates mutually stabilizes each other in plants

Since MG-132 restored both MTB and SE from 1,6-HD induced protein reduction, we next studied how protein stabilities were regulated. Upon the treatment of cycloheximide (CHX), an inhibitor of protein synthesis, MTB had a much shorter half-life in *se-1* vs Col-0, and the half-life was clearly extended in the presence of MG-132 in the two backgrounds (Fig. 4e, Extended Data Fig. 7a). Notably, the half-life of MTB in Col-0 could be slightly extended by PYR-41, an inhibitor specific for E1 enzyme for 26S proteasome pathway^16^, whereas this effect was not detected in *se-1* (Fig. 4e, Extended Data Fig. 7a). These results suggested that the IDP MTB might be subjected to destruction via 20S proteasome, and this regulation became obvious in *se-1* vs Col-0. Next, we performed cell-free protein decay assays with recombinant MTB protein. Again, GST-His-MTB was quickly degraded when incubated with the cell lysate derived from *se*-2 compared to that of Col-0, and this degradation was substantially blocked by MG-132 (Extended Data Fig. 7b,c). Furthermore, poly-ubiquitinated MTB conjugates were undetectable comparing to DMS3, a control degraded by the 26S proteasome^35^ (Fig. 4f,g). Thus, SE might stabilize MTB from 20S proteasome-mediated degradation. Further supporting this idea was that MTB could directly bind to three 20S proteasome subunits: a structural subunit, PAG1, and two proteolytically active subunits, PBE1 and PBE2 (Fig. 4h, Extended Data Fig. 7d,e).

We next performed *in vitro* 20S proteasome reconstitution assays^16^. Indeed, MTB was degraded by 20S proteasome and this process was hindered by MG-132 (Fig. 4i,k). Interestingly, pre-incubation of MTB and SE before addition of 20S proteasome could significantly stabilize both MTB and SE concordantly (Fig. 4j,l). Of note, the co-stabilization effect was also sustained by pre-incubation of LCD^FUS^-SEΔIDR1, but only weakly by SE-R718A and SEΔIDR1 (Fig. 4i–l). In line with these results, MTB protein was accumulated in *pag1* and *pbe1* vs Col-0 (Fig. 4k–m, Extended Data Fig. 7f), reminiscent of SE accumulation in 20S mutants^16^. These results indicated that SE-MTB assembly might mutually protect each other from 20S proteasome mediated degradation through co-condensation.

We next hypothesized that developmental deficit of *se* might in part result from destabilization of MTC. To test that, we generated overexpression lines of *MTA* and *MTB* in Col-0. Overexpression of *MTA* led to the accumulation of both MTB and SE (Fig. 5a). However, overexpression of *MTB* only increased in SE, but not MTA, suggesting that MTA might undergo an additional regulatory survey *in vivo* (Fig. 5a). Interestingly, once being crossed to the *se-1*; *p35S*::*Flag-MTA* could partially restore mutant developmental defects as evidenced by plant size, flowering transition, and visible enrichment of truncated SE-1 (Fig. 5a–c, Extended Data Fig. 7g). In sharp contrast, overexpression of MTB could not be achieved in *se-1*, and *se-1*; *p35S*::*Flag-MTB* plants now phenocopied *se-2*, which is a more severe allele vs *se-1* (Fig. 5a–c, Extended Data Fig. 7g). These results suggested that MTB, rather than MTA, precisely the soluble portions, displayed a SE-dependent accumulation, with a further suggestion that the *se* developmental defects were partially accredited to concurrent reduction of functional MTB.

**Figure 5.**
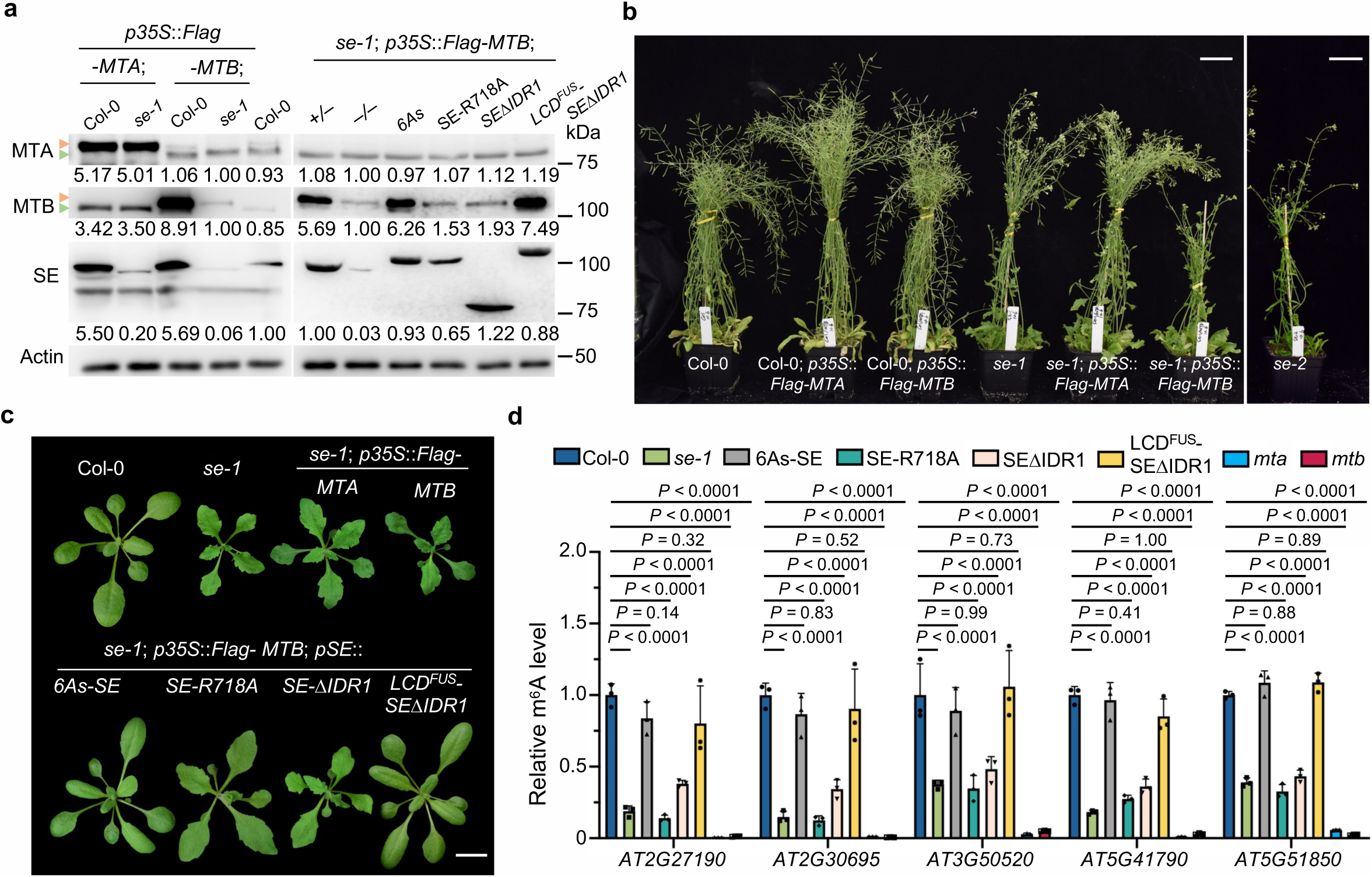
Proper phase behaviors of SE and MTB are required for protein accumulation and plant development. **a – c,** Immunoblots (**a**) and images (**b, c**) of three-week-old plants indicated the importance of both SE-MTB interaction and liquid-like properties for the plant development and protein accumulation *in vivo*. Immunoblots were performed with indicated endogenous antibodies. Actin was a loading control. Triangles indicated the Flag tagged (orange) and endogenous protein (gray) bands, respectively. Scale bars, 5 cm (**b**) and 2 cm (**c**). Three biological replicates of immunoblots were tested, at least ten independent transgenic lines were photographed, and representative images are shown. **d**, MeRIP-qPCR showed the m^6^A levels of selected genes in different complementation plants vs Col-0. Data are mean ± s.d. of three independent experiments. P values, two-way ANOVA analysis with Dunnett’s multiple comparisons test.

To further investigate if SE-MTB-regulated phenotypes entailed their interaction and condensation, we performed complementation assays of *se-1* with various SE variants. Notably, only the variants that could stabilize MTB, specifically *se-1*; *pSE*::*6As-SE* and *se-1*; *pSE*::*LCD^FUS^-SEΔIDR1*, but not *se-1*; *pSE*::*SE-R718A* and *se-1*; *pSE*::*SEΔIDR1*, could rescue the mutant phenotype (Extended Data Fig. 7h). Consistent with the morphological changes, the m^6^A modification exhibited reduced levels in the *se-1*; *pSE*::*SE-R718A* and *se-1*; *pSE*::*SEΔIDR1* hypomorphs vs WT and *se-1*; *pSE*::*6As-SE* and *se-1*; *pSE*::*LCD^FUS^-SEΔIDR1* lines, albert to a lesser extent, compared to the amount in *mta* and *mtb* mutants (Fig. 5d). Additionally, only 6As-SE and LCD^FUS^-SEΔIDR1 could accumulate MTB upon introducing into *se-1*; *p35S*::*Flag-MTB* (Fig. 5a,b). These results suggested that liquid-like phase behaviors of the SE-MTB complex could shield the proteins from 20S proteasome-mediated degradation, thereby being critical for their functionality in plants. Altogether, we conclude that SE and MTC form liquid-like co-condensates, and through this, SE maintains the solubility of MTC, promoting MTC enzymatic activity and protein stabilization.

### Methylation promotes pri-miRNA processing in *Arabidopsis*

It has been reported that m^6^A positively regulates pri-miRNA processing^22–25^, but the underlying mechanism remains unclear. Next, we hypothesized that SE-MTB interaction might impact m^6^A modification on pri-miRNAs. To test this, we first pinpointed methylation sites of pri-miRNAs. Despite of a general decline in m^6^A epi-transcriptome in *se-2*, the majority of methylation remained discernible (Fig. 1d). Taking advantage of highly accumulated defectively processed pri-miRNAs, we relaxed the peak calling stringency using a moderate threshold as log2(enrichment) > 0 and P < 0.05, and through this, 79 candidates from 127 detected pri-miRNAs were identified (Fig. 6a, Extended Data Fig. 8a, Supplementary table 5). In parallel, an independent approach^36^ using a sliding window algorithm of 25 nucleotides was applied. The m^6^A enrichment for each window was normalized using the median coverages of the corresponding genes for both IP and input. This analysis yielded 83 methylated pri-miRNAs (Supplementary table 5), with 71 shared in both algorithms (Fig. 6a). Indeed, IGV files revealed that the reads in IP samples exhibited clear enrichments that were distinct from the patterns in inputs (Fig. 6b). To further validate this, we conducted MeRIP-qPCR assays using the mixture of full-length Poly(A)+ mRNA and two standards. Consistently, pri-miRNAs detected were enriched in IP samples, but their methylation levels significantly decreased in *se-2* and *mta* compared to Col-0 (Fig. 6c). In contrast, the enrichment of methylated spikes was increased in mutants (Extended Data Fig. 8b). We further explored distribution m^6^A on pri-miRNAs and found that methylation was predominantly deposited onto the ss basal regions (Fig. 6d). Thus, SE also contributed to m^6^A methylation on pri-miRNAs mediated by MTC.

**Figure 6.**
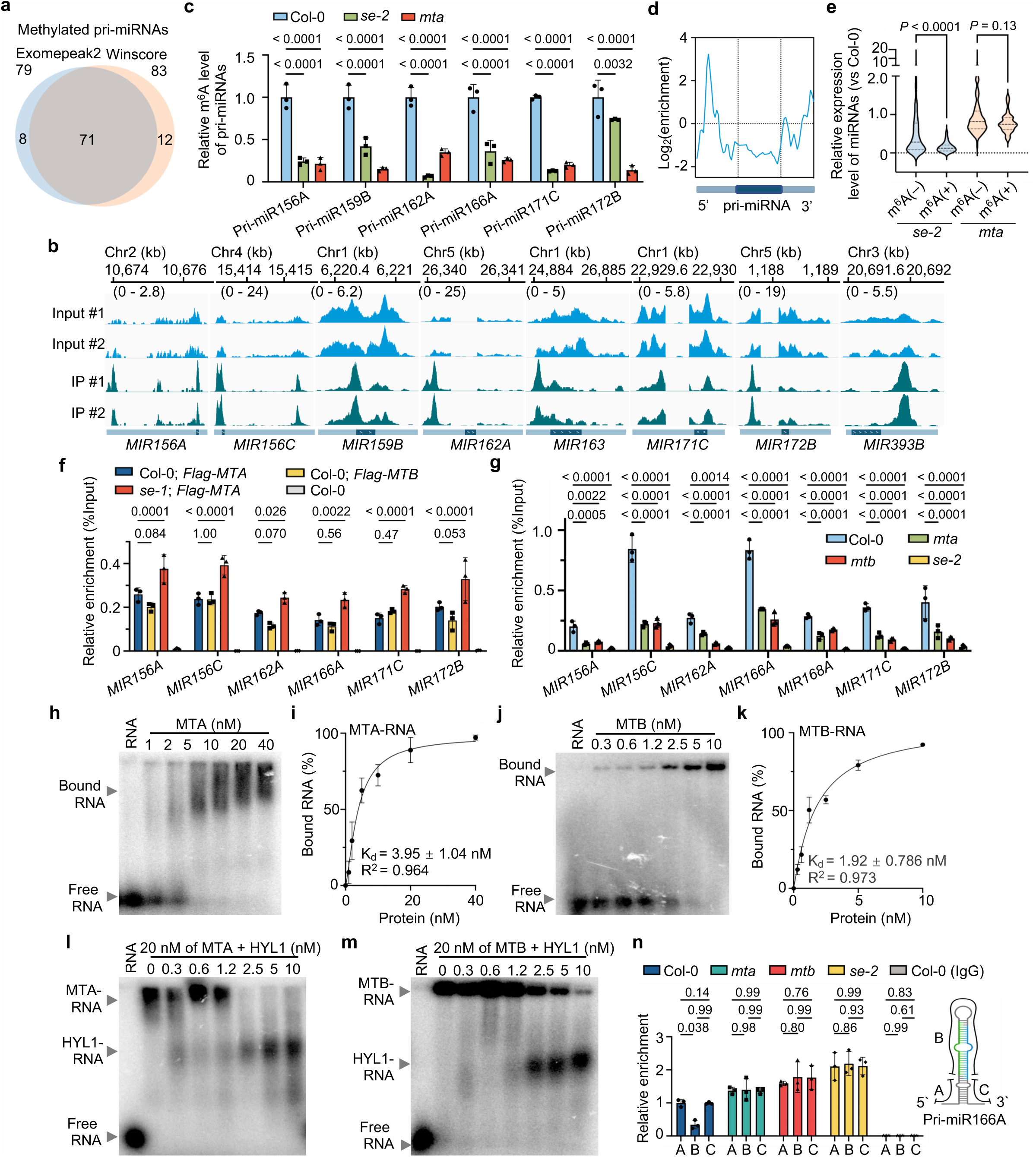
MTC facilitates miRNA biogenesis via m^6^A dependent and independent manners. **a,** Exomepeak2 and winscore algorithms identified largely overlapping methylated pri-miRNAs in *se-2*. **b,** IGV files of the selected methylated pri-miRNAs in *se-2*. The dark color region within the light horizontal line represents the pre-miRNA within the pri-miRNA. X-axis, chromosomal locations; y-axis, normalized reads. Chr, chromosome. **c,** MeRIP RT-qPCR assays validated the reduction of m^6^A levels on pri-miRNAs in *se-2* and *mta* vs Col-0. **d,** A metagene profile portrayed the average m^6^A distribution across pri-miRNAs. X-axis, relative position; y-axis, m^6^A abundance. The dark color region within the light horizontal line represents the pre-miRNA within the pri-miRNA. **e,** A violin plot illustrated that the levels of miRNAs derived from methylated pri-miRNAs were significantly reduced vs those derived from un-methylated pri-miRNAs in mutants vs Col-0. The lines represent the median and quartiles. **f, g,** ChIP-qPCR assays showed that MTA and MTB acted up-stream of SE in tested *MIRNA* loci. ChIP assays were performed with (f) anti-Flag and (g) anti-SE antibodies. **h – k,** EMSA assays (**h, j**) and the binding curves (**i, k**) showed that MTA and MTB possessed strong binding affinities to structured pri-miRNAs. **l, m,** EMSA showed that HYL1 could sequester structured pri-miRNA from MTA (**l**) and MTB (**m**). The numbers on top of gels indicated the amount of recombinant HYL1 (nM) added to 20 nM MTA and MTB. **n,** H3-RIP-qPCR assays showed increased retention of different fragments of tested pri-miRNAs along *MIRNA* loci in the mutants vs Col-0. A, B, and C refer to 5’ flanking, pre-miRNA, and 3’ flanking sequences of pri-miRNA, respectively. Data are mean ± s.d. of three independent experiments (**c**, **f**, **g**, **i**, **k**, and **n**). At least three independent experiments were performed, and representative images are shown (**h**, **j**, **l**, and **m**). P values, unpaired two-sided t-test (**e**), two-way ANOVA analysis with Dunnett’s (**c**, **f**, and **g**) and Tukey’s (**n**) multiple comparisons test. The numbers above columns are indicated p values (**c**, **f**, **g**, and **n**).

We found that the accumulation of miRNAs was significantly reduced in MTC mutants vs Col-0 (Extended Data Fig. 8c,d). The reduction was more pronounced in miRNAs derived from methylated pri-miRNAs compared to others (Fig. 6e). In lines with these results, the expression of miRNA targets was significantly increased (Extended Data Fig. 8e). The decrease in miRNA accumulation was likely attributable to a deficit in post-transcriptional processing of pri-miRNAs, as pri-miRNAs were accumulated (Extended Data Fig. 8f) whereas no obvious change of *MIR* transcription, splicing of intron-containing pri-miRNAs, or expression of key microprocessor components was detected in *mta* vs Col-0 (Extended Data Fig. 8g-j). Taken together, we pinpointed m^6^A loci on most pri-miRNAs and uncovered a positive regulatory role of writers in miRNA production.

### MTC involved in co-transcriptional processing of pri-miRNAs

Since both m^6^A deposition and the initial processing of pri-miRNAs take place co-transcriptionally, we next accessed the published SE ChIP-seq^37^ and found that SE is preferably recruited to the loci generating methylated pri-miRNAs (Extended Data Fig. 9a). Thus, we hypothesized that MTC might contribute to co-transcriptional processing of pri-miRNAs. To test this, we performed ChIP-qPCR using *MTA* isogenic transgenic plants in which the protein levels of MTC components are comparable in Col-0 and *se-1.* Results showed that MTA could be recruited to *MIRNA* loci in Col-0 and the enrichment was even increased in the absence of SE, suggestive of MTA retention onto chromatin in *se-1* vs Col-0 (Fig. 6f). Conversely, the recruitment of SE to the loci was compromised in *mta* and *mtb* (Fig. 6g), indicating that SE association with the m^6^A-conferring loci depends on MTC in this context.

To determine whether MTC could handover pri-miRNAs to microprocessor, we investigated the binding affinity of MTC to pri-miRNAs. Electrophoretic mobility shift assay (EMSA) showed a very weak RNA association with FIP37, one of MTC components (Extended Data Fig. 9b). Differently, MTA and MTB both had robust RNA binding abilities with respective K_d_ values of 3.95 (±1.92) nM (Fig. 6h,i) and 1.92 (±0.77) nM (Fig. 6j,k), but the affinities were relatively weaker than that of HYL1 which has two dsRBDs and a much stronger RNA affinity with K_d_ value of 0.74 (±0.16) nM^38^. Consistently, HYL1 could effectively absorb RNAs from MTA and MTB in competition binding assays (Fig. 6l,m). Notably, HYL1 association of pri-miRNAs is reduced in *mta*^23^, indicating that MTC might deliver pri-miRNAs to HYL1. Next, we performed Histone 3 ribonucleoprotein complex immunoprecipitation assays (H3-RIP) to examine the chromatin retention of nascent or partially processed pri-miRNAs^39^. H3-RIP-qPCR showed that the aberrant accumulation of full length pri-miRNAs rather than partially-processed intermediates was detected in H3-RIP from the MTC mutants vs Col-0 (Fig. 6n, Extended Data Fig. 9c–e). Thus, these results suggested that MTC could recruit microprocessor to *MIRNA* loci through MTB-SE interaction, and then convey pri-miRNAs to the machinery for initial processing.

### M^6^A readers affect pri-miRNA processing

Pri-miRNA processing also occurs in the nucleoplasm^39^. We found that pri-miRNAs were accumulated in the nucleoplasm of *mta* vs those of Col-0, suggestive of a critical role m^6^A in pri-miRNA processing (Fig. 7a). A series of m^6^A readers, evolutionarily conserved C-terminal region (ECT)1–3, ECT5, and cleavage and polyadenylation specificity factor 30 (CPSF30), were recovered from a proteomic assay of SE immunoprecipitates^40^ (Extended Data Fig. 10a). Among YTH-domain containing proteins, *ECT2* is the most abundantly expressed gene throughout various developmental stages and tissues (Extended Data Fig. 10b). Earlier confocal microscope imaging suggested the predominant presence of ECT2 in the cytoplasm and the observable amount in the nucleus^41,42^. To clarify this, we conducted a nucleus-cytoplasm fraction assay and found that ECT2 could indeed be detected in the nucleus fraction (Fig. 7b). Furthermore, ECT2 were readily detected in the nucleus upon the treatment of leptomycin B, a chemical that blocks the protein trafficking from nucleus to cytoplasm (Fig. 7c,d). These results suggested that ECT2 could perform some unexplored functions in nucleus. Indeed, Y2H and co-IP experiments showed that ECT2 could associate with microprocessor through SE (Fig. 7e, Extended Data Fig. 10c). Importantly the majority of tested miRNAs displayed reduced, coupled with an accumulation of some pri-miRNAs in *ect2; ect3; ect4* vs Col-0 (Fig. 7f, Extended Data Fig. 10d). These results further suggested that m^6^A might be an essential marker for effective processing of pri-miRNAs.

**Figure 7.**
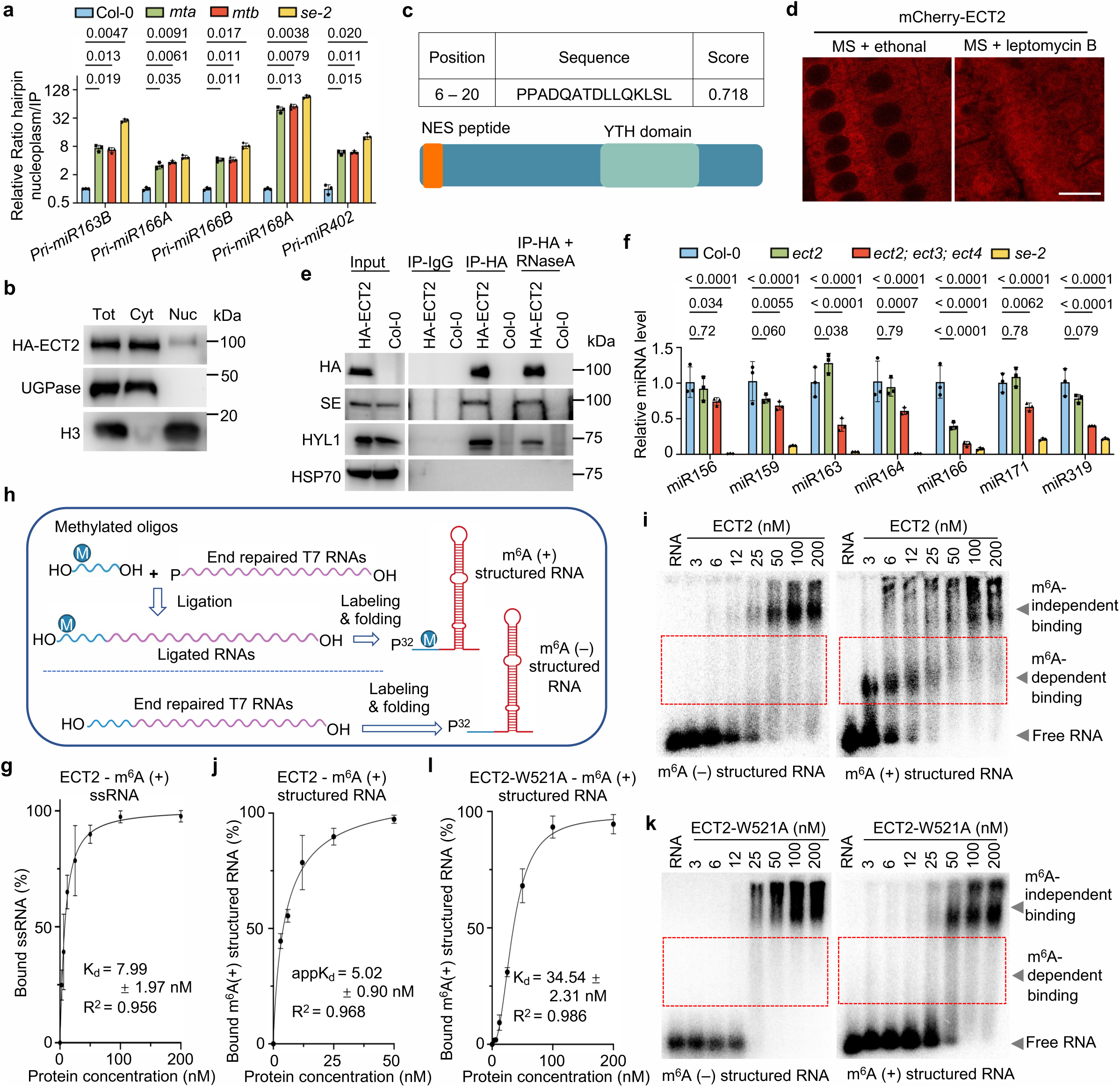
The m^6^A readers promote pri-miRNA processing. **a,** RT-qPCR assays detected accumulation of tested pri-miRNAs in the nucleoplasm fraction of m^6^A writer mutants vs Col-0. *UBQ10* was an internal control for normalization. **b,** Immunoblots of fractionation assays using three-week-old *pECT2*::*3xHA-ECT2* plants. ECT2 was detected by an anti-HA antibody. UDP-glucose pyrophosphorylase (UGPase) and Histone 3 (H3) were cytoplasmic and nuclear markers, respectively. **c**, Predicted CRM1 (exportin 1)-dependent nuclear export signal (NES) peptide in ECT2. **d**, Microscopic images showed the nuclear localization of ECT2-mCherry signals in five-day-old transgenic plants upon CRM1 inhibitor leptomycin B treatment. Ethanol was a negative control. Scale bar, 10 μm. **e**, Co-IP assays validated the interaction between ECT2 and microprocessor key components. IPs were conducted with *pECT2*::*3xHA-ECT2* plants using HA antibodies. Immunoblots were detected with indicated antibodies, respectively. HSP70 was a negative control. **f,** RT-qPCR assays showed the decreased levels of tested miRNAs in m^6^A reader mutants vs Col-0, with *se-2* serving as a control. U6 was used as an internal control for normalization. **g,** The binding curve of ECT2’s affinity to m^6^A (+) ssRNA. **h,** Schematic illustration of the strategy to synthesize m^6^A-modificated RNAs. T7 *in vitro* transcribed RNA was used for end repair, followed by ligation to a methylated oligonucleotide. The products were labeled with P^32^ and purified prior to *in vitro* folding. An unmethylated control was also synthesized using T7 *in vitro* transcription with an identical sequence. **i – l,** EMSA assays **(i, k)** and their binding curves (**j, l**) showed that ECT2 and ECT2-W521A, a variant lacking of m^6^A recognition, both possessed a binding affinity to m^6^A (–) structured pri-miRNAs and the binding affinity stimulated by the presence of m^6^A on RNAs could only be found by ECT2. Data are mean ± s.d. of three independent experiments (**a**, **f**, **g**, **j**, and **l**). At least three independent experiments were performed, and representative images are shown (**b**, **d**, **e**, **i**, and **k**). P values, two-way ANOVA analysis with Dunnett’s multiple comparisons test (**a**, **f**). The numbers above columns are indicated p values (**a**, **f**).

To further study how the m^6^A reader ECT2 contributed to pri-miRNA processing, we detected its binding affinity to different types of RNA. Purified ECT2 had a strong propensity to bind to methylated ssRNAs, but not to unmethylated ssRNAs (Fig. 7g, Extended Data Fig. 10e,f, top panels). Moreover, W521A mutation abolished ECT2 association with the methylated ssRNAs (Extended Data Fig. 10e,g, top panels), indicative of the binding’s reliance on the m^6^A group as observed previously^42^. Then, we generated structured RNAs with or without m^6^A (Fig. 7h, Extended Data Fig. 10h). Remarkably, we found that ECT2 could bind structured RNAs in a m^6^A-independent manner with a moderate K_d_ of 35.09 (± 2.34) nM (Fig. 5i, left, Extended Data Fig. 10i). However, the presence of m^6^A could booster ECT2 association with the structured RNAs with a K_d_ of 5.05 (±0.92) nM which is equivalent to or slightly lower than that of the strong RNA-binding protein HYL1^38^ (Fig. 5i, right, j). Such high affinity would enable ECT2 to recognize and pool substrates at a significantly lower concentration. On the other hand, the removal of putative ECT2 target motif UYUYU^42,43^ exhibited a slight reduction in RNA affinity (Extended Data Fig. 10f, bottom panel, j). Consistently, W521A did not impact ECT2 binding of structured RNA but erased its binding with the m^6^A-dependent portion (Fig. 7k; Extended Data Fig. 10g, bottom panel, k, l), indicating that the ECT2 association of structured RNA is independent of m^6^A modification. Together, our results indicated that ECT2 had a stronger binding affinity to m^6^A-harbored ss segments of pri-miRNAs, while also associated with the structured regions of pri-miRNAs but with a relatively weaker affinity in a m^6^A recognition-independent manner.

Finally, we assessed the impact of two-pronged RNA binding capacity of ECT2 on pri-miRNA processing. *In vitro* microprocessor reconstitution assays showed that the processing efficiency of methylated and unmethylated pri-miRNAs was comparable in the absence of ECT2 (Fig. 8a,b, Extended Data Fig. 10m), suggesting that the physical feature of m^6^A itself could not promote microprocessor activity in pri-miRNA processing. Next, we pre-incubated pri-miRNAs with a gradient dosage of ECT2 before microprocessor assays. Whereas ECT2 did not impact the processing of unmethylated pri-miRNAs, the protein could significantly enhance the cleavage efficiency of methylated pri-miRNAs (Fig. 8a,b). Furthermore, this enhancement was remarkable when the ECT2 concentrations were low but sufficient for binding the m^6^A sites. Importantly, the ECT2-mediated enhancement of the methylated pri-miRNAs relied on the recognition and binding of m^6^A by the reader ECT2 as ECT2-W521A did not enhance the cleavage of m^6^A-methylated pri-miRNAs (Fig. 8a,b). These results suggested that the m^6^A readers exemplified by ECT2 could launch on m^6^A sites located on the ss segments of pri-miRNAs while affiliating with microprocessor, consequently facilitating miRNA production. At the higher concentrations of ECT2 and its variant, the cleavage efficiency of pri-miRNAs was declined, likely because ECT2 bound and occupied the duplex regions of pri-miRNAs and blocked their processing. Collectively, we concluded that components of the m^6^A pathway play both methylation-independent and methylation-dependent roles in promoting miRNA production (Fig. 8c).

**Figure 8.**
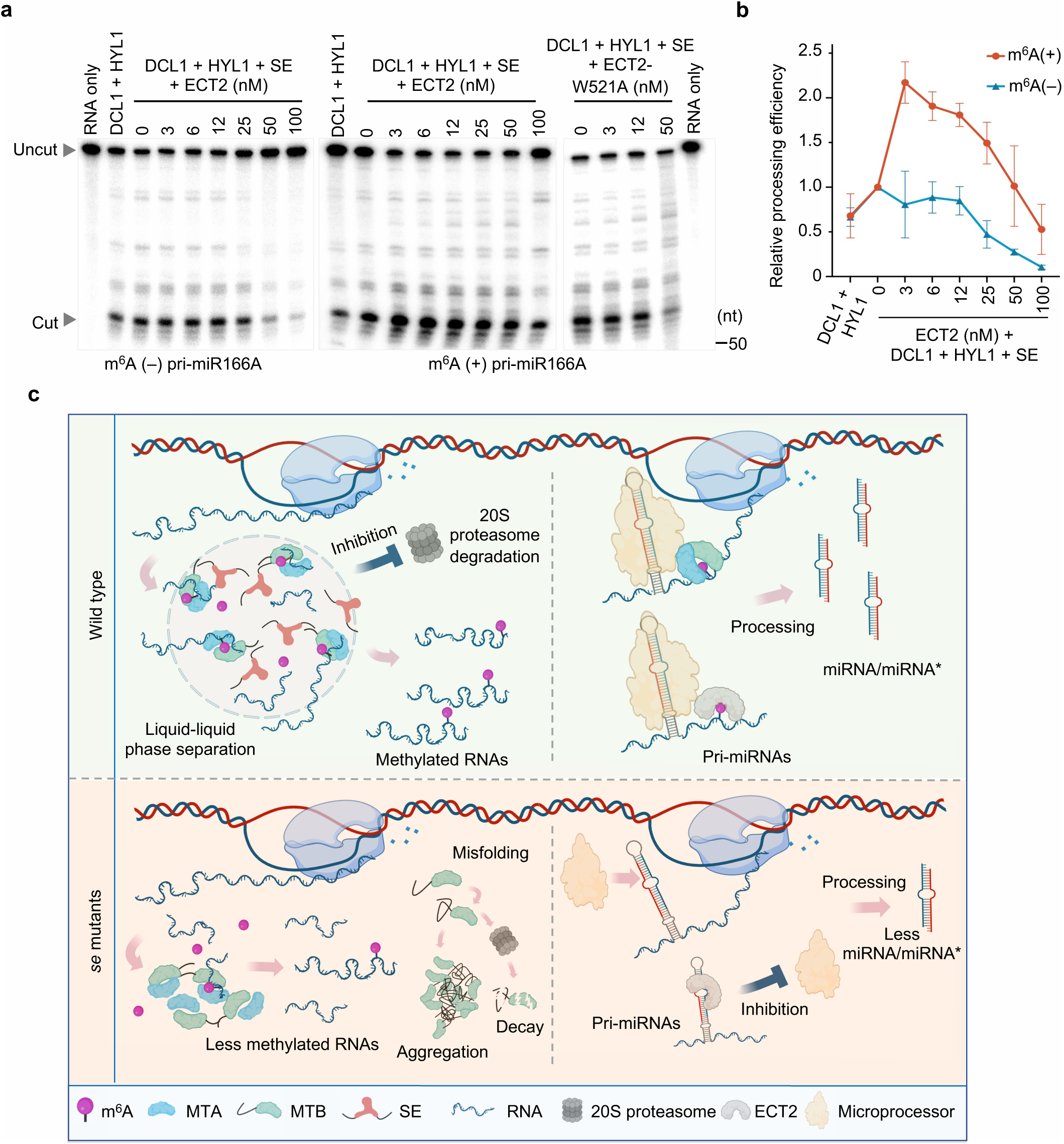
M^6^A modification promotes miRNA production in *in vitro* microprocessor reconstitution assays and a proposed model for reciprocal regulation of m^6^A modification and miRNA production machineries. **a, b,** *In vitro* microprocessor assays (**a**) and statistical analysis (**b**) of processing efficiency showed that appropriate concentrations of ECT2 were essential for effective promotion of the processing of m^6^A-harbored pri-miRNAs compared to their m^6^A-depleted counterparts. Data are mean ± s.d. of three independent experiments performed, and representative images are shown. **c,** A proposed model of cross-regulation between m^6^A modification and miRNA production machineries. Top left, MTB can co-condense with SE droplets, thereby maintaining solubility and enzymatic activity in a WT condition. Bottom left, MTB may undergo misfolding, leading to either aggregation or degradation in *se* mutants. Top right, MTC can reciprocally recruit microprocessor to *MIRNA* loci to engage in co-transcriptional processing of pri-miRNAs. Concurrently, the readers interact with both the basal or flanking region of methylated pri-miRNAs and microprocessor to facilitate processing. Bottom right, the absence of writers results in the impairment of both co-transcriptional processing and methylation of pri-miRNAs. Additionally, readers occupy structural regions within pri-miRNAs, consequently obstructing downstream processing events.

## Discussion

Here we reported that MTC and microprocessor, two otherwise distinct RNPs that are respectively engaged in epi-transcriptome modification and pri-miRNA processing, can mutually modulate each other’s function. On one side, SE acts beyond its canonical role in miRNA biogenesis and functions as a positive regulator of m^6^A modification. To do this, SE utilizes its intrinsic LLPS capacity to modulate the phase behaviors conferring the solubility of MTB, which would otherwise rapidly form insoluble condensates. The SE-driven liquid phase separation enables MTC to uptake RNA substrates inside the complex to facilitate m^6^A deposition. Meanwhile, the co-condensate can protect the IDPs from 20S proteasome mediated degradation. On the other hand, MTC promotes miRNA biogenesis by recruiting microprocessor to chromatin, initiating co-transcriptional processing of pri-miRNAs. MTC can also deposit m^6^A on ss basal and flanking regions of pri-miRNAs, granting the m^6^A readers exemplified by ECT2 to enrich the substrates and supply them to its tethered microprocessor for continuing processing.

The mechanism of SE-driven phase separation of MTB is different from the way by which SE triggers co-condensation with microprocessor or its inhibitors, SAID1/2, to regulate miRNA production^6,14,15^. The microprocessor components DCL1 and HYL1 are well-folded proteins. Without SE, these constituents are diffused but still functional, and their amount ^6,39^. even accumulates^16^. Here, MTB, an IDP, heavily relies on SE for its solubility, and stability to certain extent. In this context, SE acts an "anti-freeze" character to shift the equilibrium of MTB away from forming insoluble condensates towards the soluble condensates, conferring the MTC with robust enzymatic activity. Importantly, both SE and MTB can also stabilize each other by altering their conformations and excluding proteasome from the surrounding niche. In this perspective, SE behaves as a “chaperone”, like HSPs^44,45^, instead of a well-perceived “scaffold” for multiple RNPs^13^. Hence, co-condensation of SE and MTB represents an escape mechanism that prevents 20S-proteasome-mediated degradation of diffuse proteins, illuminating the roles of phase separation in controlling protein stability. This scenario is reminiscent of the PGL-1/-3 that could evade autophagic degradation by undergoing LLPS^46,47^.

Notably, the SE-modulated MTB phase separation is also different from previously reported condensation patterning of mammalian METTL3 or *Arabidopsis* MTA. Mammalian METTL3 and METTL14 are both structured proteins. METTL3, when fused with the IDR of CRY2, could form puncta in mammalian cell lines, while METTL14 could not^18^. In *Arabidopsis*, MTA appears to orchestrate CRY2 in a co-LLPS manner to determine m^6^A deposition of light signaling genes^19^. However, MTB does not directly interact with CRY2 and the two proteins unlikely undergo co-condensation^19^. Whether MTA-CRY and MTB-SE crosstalk to regulate light signaling pathway or other biological processes warrants further studies.

We re-discovered that m^6^A could facilitate miRNA production, and uncovered three unique features: First, we accredited the m^6^A-mediated facilitation of miRNA production to the m^6^A readers exemplified by ECT2. This is different from the mechanism in mammals where the enhancement is accredited to HNRNPA2B1, which does not contain YTH domain and is not a direct m^6^A "reader" but could influence on miRNA production via a "m^6^A switch" mechanism^25^. Second, we pinpointed the m^6^A modification sites on the ss basal and flanking regions of pri-miRNAs, clarifying earlier confusion of the possible locations of m^6^A on the stem-loop regions. We argued that the methylation in the stem-loop region that could impair pri-miRNA processing as m^6^A inherently inclines to unfold RNAs owing to the frailty of m^6^A-Uracil base pairs^25^. Furthermore, the occupation of m^6^A readers may spatially obstruct microprocessor from the accessibility to the substrates. Indeed, both plant m^6^A reader ECT2 and mammalian HNRNPA2B1^24^ could bind to the structured regions of pri-miRNAs with approximately an 8-fold diminished affinity, independent of m^6^A recognition, which might subsequently suppress the processing of substrates. In line with this, the m^6^A writers have been observed to preferably deposit m^6^A onto ssRNA regions in mammals^48^. Additionally, we elucidated that MTC could orchestrate co-transcriptional processing nearby the chromatin and post-transcriptional processing of pri-miRNAs in the nucleoplasm. Whether MTC possesses such a mechanism in other organisms awaits future investigation.

## Acknowledgements

This work was supported by grants NIH GM127414 and NSF MCB 1818082 to B. Y., NSF of Guangdong Province (Grant No. 2020B1515020007), NSFC (31771349, 32170593), and Guangdong Provincial Pearl River Talent Plan (Grant NO. 2019QN01N108) to Z.Z., and NIH R35GM151976, NSF MCB 2139857, and the Welch foundation (A-2177-20230405) to X. Z.

## Author contribution statement

X.Z. and S.Z. conceived the project and designed the experiments. Z.Z. and B.Y. independently discovered the project of SE-MTB interaction.

S.Z. performed most of the experiments.

X.L generated genetic materials and plasmids and participated in confocal microscope analysis.

Jiyun Zhu. and T.M. performed HPLC-MS and data analysis.

C.L. analyzed MeRIP-seq and sRNA-seq data.

H.B., and J.J.C. generated MTA and MTB antibodies, and validated partial results.

L.G. created lines of Flag tagged conjugated MTA and MTB overexpression transgenic plants and native promoter driven GFP tagged MTB, with either Col-0 or *se-1* background. Jiaying Zhu, T.O, and N.L. contributed to *in vitro* RNA transcription.

X.Y participated in protein purification.

H.K. and X.P. provided intellectual and experimental support.

S.Z wrote the initial draft of manuscript and X.Z. thoroughly edited the paper, all authors contributed to the proof-reading of manuscripts.

## Competing interest statement

The authors declare no competing interests.

**Correspondence and requests for materials** should be addressed to X.Z.

## Methods

No statistical methods were used to predetermine sample sizes. Randomization and blinding design were not relevant to this study.

### Plant materials and growth conditions

*Arabidopsis thaliana* ecotype Columbia (Col-0), *se-1* (CS3257), *se-2* (SAIL_44_G12), *se-3* (SALK_083196), *hyl1-2* (SALK_064863), *ect2-1* (SALK_002225C), and *ect2*; *ect3*; *ect4* (CS2110132) were used for this study.

For generating knock down lines, binary vectors including *pBA-p35S*::*amiR-MTA-a, pBA-p35S*::*amiR-MTA-b*, *pBA-p35S*::*amiR-MTB*, and *pBA-p35S*::*amiR-FIP37* were transformed into Col-0 by the floral-dip transformation method. Transgenic plants were screened for the presence of artificial miRNAs and the decrease of target transcripts using small RNA blot or RT-qPCR, respectively. Subsequently, *amiR-MTA-a* plants were used for majority of experiments as *mta*, including RNA-seq, m^6^A analysis, miRNA analysis, crossed with *pMIR167A::GUS* for GUS histochemical analysis. 20S proteosome mutants were generated in the previous study^16^.

The *pSE*::*FM-SE* and *pSE*::*gSE-eGFP* (renamed as *pSE*::*SE-GFP*) were generated in previous studies, respectively^15,38^. For generating additional complementation and mutation lines, the homozygous *se-1^-/-^*mutant was transformed with binary vectors of *pBA-pSE*::*SE* derivatives by the floral dip transformation method^49^. The transgenic plants were screened by western blot analysis and RT-qPCR. Subsequently, to test if SE and its derivatives can restore the overexpression of MTB, a background switch was achieved by crossing transgenic plants with *se-1*; *p35S::Flag-MTB*.

For generating transgenic lines, binary vectors *pH7-pMTB*::*YFP-gMTB* and *pBA-p35S*::*FM (Flag-4xMyc)-FIP37* were transformed into Col-0, and *pBA-pMTB*::*mCherry-gMTB* was dipped into *pSE*::*SE-GFP*^15^, respectively. The T1 transgenic lines expressing the tagged proteins were screened by western blot analysis or confocal microscopy. Subsequently, *pH7-pMTB*::*YFP-gMTB* was crossed with *se-1* to switch background.

Col-0 (wild-type), mutants and transgenic lines were grown on soil (Jorry Gardener/LP5) or MS plates under a 12-h light/12-h dark regime at 22° ± 1°C and ∼50% relative humidity. Typically, three-week-old plants were collected for MeRIP-seq, RNA-seq, sRNA-seq, RT-qPCR, western blot analyses, and ChIP/RIP-PCR unless specifically mentioned.

The Col-0; *pMIR167A*::*GUS*, Col-0; *p35S*::*Flag-MTA*, Col-0; *p35S*::*Flag-MTB*, *se-1*; *p35S::Flag-MTA*, and *se-1*; *p35S*::*Flag-MTB* seeds were generated by Dr. Bin Yu’s laboratory. The *ect2-1*; *ect3-1*; *ect4-2*; *pECT2*::*ECT2-mCherry* (CS2110846) and *ect2-1*; *ect3-1*; *pECT2*::*3xHA-ECT2* (CS2110850) were obtained from ABRC. The mutants were crossed with *ect2-1* to switch background(s).

### Vector construction

The plant binary constructs were made using the Gateway system (Thermo Fisher). For the majority of coding DNA sequences (CDSs) and genomic sequences, an entry vector, pENTR1A (Thermo Fisher), was used for initial cloning, followed by the LR reaction. Construction using digested vector were performed using either T4 ligations or NEBuilder^®^ HiFi DNA Assembly Master Mix with a standard procedure. The complete list of primers for all constructs can be found in Supplementary table 7.

The CDS of *MTA*, *MTB*, *N-truncated MTB* (1-634), *C-truncated MTB* (635-963), *FIP37*, ECT2, and *VIR*, and the gDNA of *gMTA* and *gMTB* were amplified and cloned into pENTR1A vector. The point mutation and truncation derivates of SE and point mutation of ECT2 were generated by PCR and cloned into a pENTR1A vector; the chimera protein of *LCD^FUS^-SEΔIDR1* was obtained with a previous approach^6^ and cloned into pENTR1A vector. The binary vector of *pBA-pSE*::*DC* and *pBA-pSE*::*mCherry-DC* was generated using 2,241 bp prior to start codon as in the previous study^38^. The same promoter sequence of *SE* was adopted to generate *pBA-pSE*::*mCherry-DC*. LR reaction was performed with *pBA-pSE*::*DC* and pENTR1A carrying different variants, yielding *pBA-pSE*:*:SE* and its derivates.

To generate native promoter driven plasmids, 3,223 bp and 2,374 bp before start codon were chosen as promoter for *MTA* and *MTB*, respectively. The promoters were amplified and cloned into pBA002a-DC-CFP (digested by BamH I and Xba I), *pBA002a-mCherry-DC* (digested by BamH I and Xba I), and pH7WGY (digested by SpeI and Sac I) through HIFI recombination (NEB), yielding *pBA-pMTA*::*DC-CFP*, *pBA-pMTB*::*mCherry-DC*, and *pH7-pMTB*::*YFP-DC*. For protein transient expression in *Arabidopsis* mesophyll protoplast and tobacco plants, *pENTR1A-MTA, pENTR1A-MTB*, and *pENTR1A-SE* and its variants were used in LR reaction to generate *pBA-p35S*::*MTA-CFP*, *pBA-pMTA*::*MTA-CFP*, *pH7-p35S*::*YFP-MTB*, *pH7-pMTB*::*YFP-MTB*, *pBA-p35S*::*mCherry-SE*, *pBA-pSE*::*mCherry-SE* and its derivates. To make transgenic plants, *pENTR1A-gMTB* was used in LR reaction to obtain the binary construct of *pBA-pMTB*::*mCherry-gMTB* and *pH7-pMTB*::*YFP-gMTB*.

For generating artificial miRNA constructs, a template of *pENTR-amiR159*^16^ was amplified with specific primers and the PCR products were subjected to Xma I and Bgl II digestion. The resulting fragments were ligated to Xma I and Bgl II digested *pENTR/D-amiR-PAG1*^16^ backbone to generate *pENTR/D-amiR-gene*. For generating Y2H constructs, the destination vectors *pGADT7-DC* and *pGBKT7-DC* were used in LR reaction. For generating BiFC constructs, *pBA-p35S*::*nYFP-DC* and *pBA-p35S*::*cYFP-DC* were used in LR reaction. For generating LCI constructs, *pCambia1300-p35S*::*cLUC-HA-DC* and *pCambia1300-p35S*::*MYC-DC-nLUC* were used in LR reaction. Plasmids containing microprocessor components and 20S proteasome subunits used in Y2H, BiFC, and LCI assays were generated in previous study^16^.

For protein purification constructs, CDSs of *MTA*, *FIP37*, *ECT2, ECT2-W521A*, *SE* and its derivates were cloned into pET28a-6xHis-SUMO vector through BamH I and Sal I restriction sites. *MTB* was cloned into Xho I / Sma I-digested pAcGHLT-C vector. For fluorescence tagged constructs, *mCherry-FUS^LCD^*, *MTA-CFP*, *YFP-MTB*, *mCherry-SE*, and *mCherry-SE-variants* were amplified from *pBA-pSE*::*mCherry-LCD^FUS^-SEΔIDR1*, *pBA-p35S*::*MTA-CFP*, *pH7-p35S*::*YFP-MTB*, and *pBA-p35S*::*mCherry-SE* and its derivates, respectively.

### Leptomycin B treatment

Five-day-old vertical cultured seedlings were transferred to 3 mL liquid MS-medium for 12 hours, then adding leptomycin B (Enzo Life Sciences) to final 2.5 μM or equal amount of 100% ethanol to medium and incubated for another 12 hours.

### Nuclear-Cytoplasmic Fractionation

The three-week-old plants were used for nuclear-cytoplasmic fractionation. Briefly, 0.5 grams of plant was homogenized with liquid nitrogen to obtain fine powder and then suspended with 6 mL of lysis buffer (20 mM Tris-HCl, pH 7.5, 20 mM KCl, 2 mM EDTA, 2.5 mM MgCl_2_, 25% glycerol, 250 mM sucrose, 5 mM DTT, 50 μM MG-132 and 1× protease inhibitor (Roche)). 100 μL of lysate was collected as “total”. The lysates were then filtered through two layers of Miracloth twice and centrifuged at 1,500 x g for 10 minutes at 4°C. After centrifugation, 1 mL of the supernatant was collected as “cytoplasmic fraction” followed by another centrifuged at 10,000 x g for 10 minutes at 4 °C. The pellet from 6 mL lyaste was washed with 5 mL of nuclear resuspension buffer 1 twice (20 mM Tris-HCl, pH 7.4, 25% glycerol, 2.5 mM MgCl_2_ and 0.2% Triton X-100). After washing, the pellet was resuspended with 500 μL of nuclear resuspension buffer 2 (20 mM Tris-HCl, pH 7.5, 0.25 M sucrose, 10 mM MgCl_2_, 0.5% Triton X-100, 5 mM β-mercaptoethanol, 50 μM MG-132 and 1× protease inhibitor (Roche)).Then, the resuspended sample was carefully added on the top of 500 μL of nuclear resuspension buffer 3 (20 mM Tris-HCl, pH 7.5, 1.7 M sucrose, 10 mM MgCl2, 0.5% Triton X-100, 5 mM β-mercaptoethanol, 50 μM MG-132 and 1× protease inhibitor (Roche)), and they were centrifuged at 16,000 x g for 45 minutes at 4°C. The final pellet was resuspended in 300 μL of nuclei sonication buffer (40 mM Tris-HCl pH 7.4, 150 mM NaCl, 5 mM MgCl_2_, 1% SDS, 1 mM DTT, 1% Triton X-100, 2% glycerol, 1 mM PMSF, 50 μM MG-132 and 1× protease inhibitor (Roche)). Sonication was performed with 15 cycles of 30-second sonication and 90-second pause. Following centrifugation at 21,000 x g for 15 minutes at 4°C, the supernatant was collected as “nuclear fraction”. Makers was adopted from previous study^16^

### Nucleoplasmic RNA extraction

Nucleoplasmic fraction was firstly obtained with a modified method adopted from previous study^50^. Briefly, two grams of three-week-old seedlings was homogenized with liquid nitrogen to obtain fine powder and then suspended with 12 mL Honda buffer (0.44 M sucrose, 1.25% (w/v) Ficoll, 2.5% (w/v) dextran T40, 20 mM HEPES-KOH pH 7.4, 10 mM MgCl2, 1 mM dithiothreitol (DTT), 1× protease inhibitor (Roche) and 10 U/mL SUPERase-In RNase Inhibitor). Debris was removed by passing through two layers of Miracloth twice. The pellet was obtained by centrifugation at 1,500 x *g* for 10 minutes at 4°C, followed by wash with 5 mL of Honda buffer containing 0.5% Triton X-100 for three times. The final pellet was collected as nuclei. The nuclei were then fully resuspended in 1 mL of nuclear lysis buffer (25 mM HEPES-KOH pH 7.4, 300 mM KCl, 0.8 M urea, 1 mM DTT, 1% Tween-20, 20 U SUPERase-In RNase Inhibitor) and centrifuged at 21,000 x *g* for 10 minutes at 4°C. The supernatant was collected as the nucleoplasmic fraction. The nucleoplasmic fraction was then subjected to TRIzol treatment for RNA isolation,

### Sedimentation assay

Ten-day-old seedling and three-week-old plants were collected and frozen in liquid nitrogen. Samples were ground into a fine powder and aliquoted into Eppendorf tubes. To extract total protein, a 5-fold weight volume of 1x SDS-loading buffer was added to the powder. Alternatively, the powder was mixed with four-fold weight volume of lysis buffers (40 mM Tris-HCl pH 8.0, 50 – 800 mM NaCl, 2 mM MgCl_2_, 50 μM ZnCl_2_, 1 mM DTT, 0.5% Triton X-100, 2% glycerol, 1 mM phenylmethyl sulfonyl fluoride (PMSF), 50 μM MG-132, and 1 x Complete EDTA-free protease inhibitor (Roche)) and rotated for 20 minutes at 4°C. Subsequently, the lysate was subjected to centrifugation at 21,000 x g for 15 minutes at 4°C. The resulting supernatant was collected and mixed with 1-fold weight volume of 5x SDS loading buffer, referring to soluble fraction. Meanwhile, the resulting pellet was resuspended 5-fold volume of 1x SDS loading buffer and collected as insoluble fraction.

To quantify, the enrichment was calculated by ((Supernatant^Protein^ / Supernatant^Control^)/(Pellet^Protein^ / pellet^Control^)). The relative enrichment was further compared with that of Col-0 at the condition of 50 mM NaCl lysis buffer, in which the value was arbitrarily set as 1.

### Western blot

The blots were detected with primary antibodies against Flag (Sigma-Aldrich, A8592, IB 1:5000), actin (Sigma-Aldrich, A0480, IB 1:5000), histone 3 (Agrisera, AS10 710, IB 1:5000), AGO1 (Agrisera, AS09 527, IB 1:1000), SE (Homemade, IB 1:5000)^16^, ubiquitin (Santa Cruz Biotechnology, sc8017, IB 1:1000), HYL1 (homemade, IB 1:5000)^16^, YFP & GFP (Roche, 11814460001, IB 1:2000), DCL1 (Agrisera, AS12 2102, IB 1:1000), His (Sigma-Aldrich, H1029, IB 1:5000), HSP70 (Agrisera, AS08 371, IB 1:5000), UGPase (Agrisera, AS14 2813, IB 1:5000), MTA (homemade, IB 1:1000), and MTB (homemade, IB 1:1000). The control IPs were conducted with IgG (Sigma-Aldrich, I4506, 3 μg per IP). Secondary antibodies were goat-developed anti-rabbit (Cytiva, NA934, IB 1:3000) and anti-mouse IgG (Cytiva, NA931, IB 1:3000). The anti-MTA and anti-MTB antibodies were generated by Dr. Zhonghui Zhang’s laboratory using full-length recombinant proteins from *E. coli*. Detailed validation is described in Reporting Summary and shown in Supplementary Data. 1.

### Co-immunoprecipitation

Three-week-old Arabidopsis thaliana plants were collected and immediately frozen in liquid nitrogen before being ground into a fine powder. The total protein was extracted with the buffer containing 40 mM Tris-HCl pH 8.0, 150 mM NaCl, 2 mM MgCl2, 50 μM ZnCl_2_, 1 mM DTT, 0.5% Triton X-100, 2% glycerol, 1 mM phenylmethyl sulfonyl fluoride (PMSF), 50 μM MG-132, and 1 pellet per 10 mL of Complete EDTA-free protease inhibitor (Roche). The supernatant obtained after centrifugation was incubated with antibody-conjugated beads at 4°C for 2 h. Anti-Flag M2 magnetic beads (Sigma-Aldrich, M8823) and GFP-trap magnetic beads (ChromoTek, gtma) were used for enriching Flag-and YFP-tagged proteins, respectively. In contrast, protein A/G beads conjugated with indicated antibodies were employed for other targets. Immunoprecipitate with IgG was used as a negative control.

Unspecific-bound binding was eliminated by five consecutive washes with IP buffer, and the beads were subsequently boiled with 2× SDS-loading buffer for western blot analyses. For the co-IP assays with RNase digestion, 50 μg/mL of RNase A (Sigma-Aldrich, R4875) was added to lysates and then incubated at room temperature for 10 minutes before immunoprecipitation.

### RNA immunoprecipitation (RIP) / Chromatin immunoprecipitation (ChIP) assay

Three-week-old seedlings were cross-linked using the buffer (20 mM Tris-HCl pH 7.4, 1 mM PMSF, 1 mM EDTA, 0.4 M sucrose, 1% formaldehyde) and ground to fine powder in liquid nitrogen. Plant nuclei were isolated from two-gram materials and homogenized with 300 μL of the nuclei sonication buffer (40 mM Tris-HCl pH 7.4, 150 mM NaCl, 5 mM MgCl_2_, 50 μM ZnCl_2_, 1% SDS, 1 mM DTT, 1% Triton X-100, 2% glycerol, 1 mM PMSF, 50 μM MG-132, and 1 pellet per 10 mL Complete EDTA-free protease inhibitor, and 10 U SUPERase-In RNase Inhibitor). Sonication was performed with 15 cycles of 30-second sonication and 90-second pause for MTA immunoprecipitation, and 12 cycles for H3 immunoprecipitation^51^. Following centrifugation at 21,000 x g for 15 minutes at 4°C, the supernatant was diluted with 9-fold volume of dilution buffer (40 mM Tris-HCl pH 7.4, 150 mM NaCl, 5 mM MgCl2, 50 μM ZnCl_2_, 1 mM DTT, 2% glycerol, 1 mM PMSF, 50 μM MG-132, 1 pellet per 10 mL Complete EDTA-free protease inhibitor, and 10 U/mL SUPERase-In RNase Inhibitor) prior to immunoprecipitation. The immunoprecipitations were performed at 4°C with agitation for 2 h (RIP) or overnight (ChIP). Either endogenous antibodies of MTA and H3 or Flag antibodies were used for RIP or ChIP, respectively. IP samples were subjected to a series of washing steps involving dilution buffer, high salt wash buffer (20 mM Tris-HCl pH 7.4, 500 mM NaCl, 5 mM MgCl_2_, 50 μM ZnCl_2_, 1 mM DTT, 1 mM PMSF, 0.5% Triton X-100, 50 μM MG-132, 1 pellet per 10 mL Complete EDTA-free protease inhibitor, and 10 U/mL SUPERase-In RNase Inhibitor), LiCl wash buffer (0.25 M LiCl, 1% NP-40, 1% sodium deoxycholate, 20 mM Tris-HCl pH 7.4, 1 mM EDTA) and Proteinase K buffer (20 mM Tris-HCl pH 7.4, 200 mM NaCl, and 10 U SUPERase-In RNase Inhibitor). Each washing step included a quick buffer resuspension and centrifugation at 1,000 x *g* for 1 minute at 4°C followed by a second wash with 5 minutes agitation. RNA or DNA was eluted from de-cross-linking with Proteinase K solution (20 mM Tris-HCl pH 7.4, 200 mM NaCl, 1 mg/mL Proteinase K, 1% SDS, and 10 U SUPERase-In RNase Inhibitor) on a Thermomixer shaker at 65°C for 6 h. The RNA was then purified using RNA Clean & Concentrator kits (ZYMO, R1017) and treated with DNase before RT-qPCR analysis. The DNA was subjected to RNase treatment and then purified using DNA Clean & Concentrator kits (ZYMO, D4004).

### Real-time quantitative polymerase chain reaction (RT-qPCR)

DNA was removed before reverse transcription with a mixture of random hexamers and oligo(dT)_12-18_. Primers for stem-loop RT-qPCR of miRNA expression can be found in the previous study^52^. Relative expression level was calculated with the 2^−ΔΔCt^ method. Primers were listed in Supplemental Table 7.

### sRNA sequencing and bioinformatics analysis

Total RNA was extracted from three-week-old plants, followed by separation of small RNAs in 15% denaturing urea gels and purification for library generation. sRNA was separated in 15% denaturing urea gels and purified for library generation. The library was prepared according to a homemade method^53^, which includes adaptor ligation, reverse transcription, PCR amplification, and purification. The resulting library was then subjected to high-throughput sequencing, and bioinformatics analysis was performed to generate reads per million (RPM) values based on mapped reads, and miR161.1, as it consistently showed no changes between *mta* and Col-0 plants in both our sRNA RT-qPCR and published sRNA sequencing assays^23^, was used for normalization.

### Yeast-two-hybrid (Y2H) assays

The positive control was adopted from previously study^54^. Briefly, different combinations of constructions were co-transformed into the yeast strain AH109 and grew at SD plates with a screening condition. At least 10 independent colonies were tested for each interaction.

### In vitro 20S proteasome decay assay

The 20S proteasome and the control IP were purified from five-gram-powder of *pPAG1*::*gPAG1-FM* transgenic plants^16,55^ and Col-0 using 300 μL of M2 bead, respectively. The immunoprecipitates were subjected to stringent washes with 800 mM NaCl buffer. The purified proteasome tested active using succinyl-Leu-Leu-Val-Tyr-7-amido-4-methylcoumarin (Suc-LLVY-AMC) (Sigma-Aldrich, S6510) was used for further assay^55^. The concentrations of 20S proteasome and test proteins were determined by the Bradford method using bovine serum albumin as a standard. Specifically, recombinant purified SE (100 nM) and Flag-MTB (10 nM), immunoprecipitated from *p35S*::*Flag-MTB* plants, were incubated with purified 20S proteasome (20 nM) in a reaction buffer (50 mM Tris-HCl pH 7.5, 150 mM NaCl) containing 2% DMSO or 50 μM MG-132. The aliquots were incubated in a PCR machine at 22°C with 37°C lid. The reaction was stopped by adding SDS–PAGE loading and boiling.

### Expression and purification of recombinant protein

GST-6×His-DCL1 and HYL1 proteins were purified in the previous work^38^. For purification of SE and its derivatives fused with or without mCherry, MTA, MTA-CFP, FIP37, LCD of FUS, ECT2 and ECT2-W521A, recombinant proteins were purified in bacterial cells. To induce recombinant protein expression in *E. coli*., transformed into BL21(DE3) cells were grown in Luria Broth (LB) medium at 37°C until OD_600_ = 0.6. Recombinant protein expression was induced by adding isopropyl-β-d-thiogalactopyranoside (IPTG) to final 0.5 mM and incubated at 16°C overnight. The induced cells were collected and re-suspended in the lysis buffer (20 mM Tris-HCl pH 8.0, 800 mM NaCl, 2% glycerol, 1 mM β-mercaptoethanol, 1 mM PMSF, 0.5% Triton X-100, 1 pellet per 50 mL Complete EDTA-free protease inhibitor (Roche)) and disrupted by the high-pressure homogenizer (Microfluidics). The cleared lysate was supplemented with 20 mM imidazole and loaded on a HisTrap HP column (GE Healthcare, Cat#: 17-5248-02). The peak fractions containing the recombinant proteins were pooled and digested by adding 30 μg of His-SUMO Protease during dialysis with 2 L buffer (20 mM Tris-HCl pH 8.0, 500 mM NaCl, 1 mM β-mercaptoethanol) at 4°C for 12 h. Ni-beads were used to remove uncut proteins and His-SUMO protease. Cleaved recombinant proteins were loaded onto a HiLoad 16/600 Superdex 200 pg column for further purification with an identical condition (20 mM Tris-HCl pH 8.0, 500 mM NaCl). The two-step purified proteins were either concentrated or dialyzed with 2 L dialysis buffer (20 mM Tris-HCl pH 8.0, 150 mM NaCl, 5 mM β-mercaptoethanol, 40% glycerol) at 4°C for 12 hours. Concentrated proteins were immediately used for in vitro phase separation assays. The dialyzed proteins were flash-frozen in liquid nitrogen and then stored at −80°C.

For purification of MTB and YFP-MTB, *pAcGHLT-GST-6xHis-MTB* and *pAcGHLT-GST-6xHis-YFP-MTB* were co-transfected with BaculoGold baculovirus DNA (BD Biosciences, Cat # 554740) into sf9 insect cells (BD Biosciences Cat# 554738; authenticated by the vendor BD Biosciences) to generate recombinant baculovirus according to the manufacturer’s instructions. The recombinant viruses were amplified for two rounds, and P3 virus was collected for large-scale protein expression. P3 virus was added to 2.5 × 10^6^ sf9 insect cells per mL for propagation, and insect cells were collected 70 h later. The cell pellet was re-suspended in lysis buffer (50 mM Tris-HCl pH 8.0, 1 M NaCl, 2% glycerol, 2 mM β-mercaptoethanol, 3 mM PMSF, 1% Triton X-100, 1 pellet per 50 mL Complete EDTA-free protease inhibitor (Roche)) and disrupted with a high-pressure homogenizer (Microfluidics). The lysate was load onto to HIS-trap HP column and the elutes containing the recombinant proteins were pooled and subjected to two continued dialysis with 1 L buffer (20 mM Tris-HCl pH 8.0, 500 mM NaCl, 1 mM β-mercaptoethanol) at 4°C for 2h per time. Then, the proteins were loaded onto GST-trap and the elutes containing the recombinant proteins were pooled and subjected to thrombin protease (Sigma) digestion ith 2 L buffer (20 mM Tris-HCl pH 8.0, 500 mM NaCl, 1 mM β-mercaptoethanol) at 4°C for 16 h. Cleaved recombinant proteins were loaded onto a HiLoad 16/600 Superdex 200 pg column for further purification with an identical condition (20 mM Tris-HCl pH 8.0, 500 mM NaCl). Purified proteins were either concentrated for in vitro phase separation assays or dialyzed with 2 L buffer (20 mM Tris-HCl pH 8.0, 500 mM NaCl, 5 mM β-mercaptoethanol, 40% glycerol) at 4°C for 6 h for storage at −80°C.

### In vitro condensate formation

Aliquots of tested proteins were centrifuged at 15,000 × *g* for 5 minutes to remove any aggregates. Droplet formation was induced by diluting the proteins using Tris-HCl pH 7.5, 20 mM. All of *in vitro* phase separation experiments were carried out in the buffer with final salt concentration of 20 mM Tris-HCl pH 7.5, 150 mM NaCl, 50 μM ZnCl_2_ without crowding agent, unless mentioned in figure legends. Slides were imaged at a 5-minute interval starting from the preparation to allow ample time for droplet settling on the glass surface.

We conducted in vitro co-condensate formation through three distinct approaches in Figure 2c. In the first method, as depicted in the top panel of Figure 2c, individual proteins were initially adjusted to concentrations ranging from 800 mM to 150 mM in the protein buffer before their combination. Subsequently, the resultant mixture was examined using confocal microscopy. In the second approach, as illustrated in the middle panel of Figure 2c, different proteins were initially combined in a solution containing 800 mM NaCl. The salt concentration was then reduced to 150 mM through dilution before further analysis. Finally, as depicted in the bottom panel of Figure 2c, SE droplets were first formed prior to the sequential addition of other proteins and RNAs.

### Confocal microscopy and analysis

All image acquisitions were carried out using a Leica stellaris 5 laser-scanning confocal microscope with a HC PL APO 63x /1.40 oil-immersion objective and LAS X Life Science Microscope Software. Sequential scanning module was applied when two or more fluorescence signals expected.

For observation of in vitro condensates, the excitation wavelengths of CFP, GFP, YFP, mCherry, and Cy5 are 405 nm, 488 nm, 514 nm, 587 nm, and 651nm, respectively. The corresponding emissions for each of these proteins are 470 – 515 nm, 500 – 530 nm, 525 – 550 nm, 597 – 630 nm, and 661 – 700 nm, respectively. For observation of protoplast, root tip and pollen samples, the excitation wavelengths of CFP, GFP, YFP, and mCherry were 405 nm, 488 nm, 514 nm, and 587 nm, respectively. Accordingly, the emissions for each of these proteins are 470 – 515 nm, 498 – 530 nm, 524 – 550 nm, and 597 – 630 nm, respectively. Chlorophyll auto-fluorescence was detected with 680 – 710 nm.

To perform time-lapse microscopy of *in vitro* condensate fusion, the samples were observed as described method without a 15-minute interval, and images were captured every 5 second over a duration of 15 minutes. Time-lapse microscopy of *in vivo* condensate fusion was executed by acquiring images every 5 second over a duration of 5 minutes.

To perform 3D rendering analysis of *in vitro* and *in vivo* assays, condensates on the glass surface and the signals located in the same layer as the cell nucleus were scanned with LIGHTNING module, respectively. 3D stimulation was carried out with LAS X Life Science Microscope Software.

### Bimolecular fluorescence complementation assay

The isolation and transfection of *Arabidopsis* mesophyll protoplasts from four-week-old Col-0 and *se-1* plants were used. Approximately 12 hours after transfection, the fluorescence signals in the protoplasts were visualized using Leica stellaris 5 laser-scanning confocal microscope.

### Fluorescence recovery after photobleaching

All FRAP assays were carried out using Leica stellaris 5 laser-scanning confocal microscope with a HC PL APO 63x /1.40 oil-immersion objective and LAS X Life Science Microscope Software.

Condensates were subjected to photobleaching using a laser intensity of 100% at corresponding emission, and pre-bleaching fluorescence intensity was obtained by capturing five images. Fluorescence intensity was then recorded every 5 seconds for 10 minutes after bleaching. FRAP analysis was performed using fluorescence intensity values and double normalization with pre-bleaching and un-bleaching regions, as well as dynamic range scaling from 0 to 1.0.

### Liquid chromatography-mass spectrometry

Isolation of mRNA from total RNAs was performed using Olig(dT)_25_ Dynabeads (NEB, S1419S). Next, 500 ng of mRNAs were subjected to enzymatic digestion with Nuclease P1 (NEB, M0660S) at 37°C for 6 h, followed by the addition of NH_4_HCO_3_ to a final concentration of 110 mM and 1 U of alkaline phosphatase (Sigma-Aldrich, P6774), and incubation at 37°C for another 6 h. Enzyme activity was quenched by heating samples at 70°C for 10 minutes, then be placed on ice for 5 minutes. After centrifugation at 21,000 x *g* for 15 minutes, the supernatants were collected for further analysis. Pure commercial adenosine (Sigma-Aldrich, A1852) and m^6^A (TargetMol, T6599) standards were also subjected to the same procedure, and standard curves were generated by running the standards of a series of concentration.

High-performance liquid chromatography (HPLC) analysis was carried out using an UltiMate 3000 HPLC system equipped with a Phenomenex Luna 5 μm C18(2) 100 Å, 4.6 mm × 50 mm column coupled to MS analysis by an ISQ EM single quadrupole mass spectrometer. The mobile phase consisted of buffer A (0.1% formic acid in water) and buffer B (100% acetonitrile). Nucleosides were quantified using the nucleoside-to-base ion mass transitions of m/z 268.0 (A) and m/z 282.0 (m^6^A). The efficiency of the system was tested using commercial methylated and unmethylated standards.

### Luciferase complementation imagine

All of the constructs were then transformed into Agrobacterium tumefaciens strain ABI. The OD values were monitored using an ultraviolet-visible spectrophotometer, then the constructs were co-infiltrated with equal ratio into combinations into *N. benthamiana*. The signal of luciferase complementation was captured 72 hours after inoculation.

### In vitro transcription and labelling of transcripts

In vitro transcription and 5’ labelling of RNA substrates was performed as described^56^. RNA substrates were transcribed under T7 promoter *in vitro* using PCR products or annealed oligos as templates, followed by DNase treatment. The RNAs were fractionated with denaturing Urea-PAGE gels and extracted from gel slices for further analysis. Structured RNAs were obtained by heating to 95°C for 3 minutes in RNA folding buffer (40 mM Tris-HCl pH 7.4 and 50 mM KCl) and gradually cooled down to room temperature. Primers and oligos sequences are listed in Supplementary Table 7.

For internal labeling of RNA, a mixture of 10 mM Cy5-UTP (Enzo Life Science) and 50 mM UTP mixture was used for *in vitro* transcription.

For 5’-end labeling of RNA, 5 pmol of dephosphorylated RNAs were labeled with ^32^P-γ-ATP (PerkinElmer) by T4 Polynucleotide Kinase (New England Biolabs). Purified RNAs were adjusted to 1000 counts per minute (c.p.m.)/μL.

### Methylated pri-miRNA synthesis

Methylated pri-miRNAs were obtained by ligating a methylated oligo and an unmethylated RNA. In the RNA ligation reaction, a donor RNA of extended length was obtained from in vitro transcription, and then used for dephosphorylation followed a T4 Kinase treatment to obtain a monophosphate group. A shorter and methylated acceptor RNA was commercially synthesized with hydroxyl groups at both ends. Subsequently, the donor and acceptor RNAs were mixed at 1:10 by molecular weight for ligating reaction (1x RNA ligase 1 buffer, 5% of DMSO, 1 mM of ATP, 1 U of SUPERase-In RNase Inhibitor, 20% of PEG8000, 1 μL of RNA ligase 1) at room temperature for 5 h. RNACleanup XP beads (Beckman Coulter, A63987) were then used to recover RNA with a length greater than 100 nt, which was subsequently subjected for 5'end ^32^P-γ-ATP labeling and purification through denaturing urea gel. Finally, the purified RNAs is folded as described above and adjusted to 1000 c.p.m/μL.

### Electrophoretic mobility shift assays

Buffers used in EMSA assays were adopted from previous study^38^. Briefly, recombinant proteins and labelled RNA were mixed in the EMSA buffer (20 mM Tris-HCl pH 8.0, 95 mM NaCl, 1 mM DTT, 0.1% NP-40, 1 U/μl SUPERase-In RNase Inhibitor). The final pooled concentration of NaCl was ∼150 mM, of which 50 mM was from protein solution and 5 mM from RNA solution. Mixtures were incubated at room temperature for 30 minutes and resolved on native 1% agarose gel. The images were quantified with ImageJ. The K_d_ and apparent K_d_ were calculated using Prism 9 (GraphPad) software fitting a Hill slope model.

### *In vitro* microprocessor reconstitution assay

In vitro Microprocessor dicing activity assays were performed according to the established protocol^38^. RNA substrate with 500 ∼ 2,000 c.p.m signal intensity was subjected to incubation with ECT2, ECT2-W521A, and His-sumo at indicated molecular amounts at room temperature for 10 minutes. The reaction was further initiated by adding 2 pmol of SE, followed by a second incubation of 10 minutes. A protein mixture containing 2 pmol of HYL1 and 1 pmol of DCL1 was subsequently added to the RNA-proteins mixture, and the final pooled salt concentration was adjusted to 20 mM Tris-HCl pH 7.5, 50 mM KCl, 4 mM MgCl_2_, 1 mM DTT, 5 mM ATP, 1 mM GTP, and 1 U/μl SUPERase-In RNase Inhibitor to initiate processing at 37°C for 20 min. The reaction was stopped by adding an equal volume of 2x denaturing loading buffer and boiling the mixture at 95°C for 10 min. The processed products were separated using a denaturing urea gel electrophoresis. Quantification of the Microprocessor dicing cleavage efficiency was carried out by the ratio of processed products comparing to unprocessed pri-miRNAs.

### Mesophyll protoplast assay

Protoplasts were prepared from *Arabidopsis* mesophyll cells. Four-week-old *Arabidopsis* leaves were meticulously sliced into slender strips and subjected to enzymatic digestion in a solution (1.25% cellulose, 0.3% macerozyme, 0.4 M mannitol, 20 mM KCl, 20 mM MES, 10 mM CaCl_2_, 5 mM β-mercaptoethanol, and 0.1% BSA). Vacuum infiltrate leaf strips for 30 minutes, then incubated in darkness for 3 hrs at room temperature until the protoplasts were released. Following filtration through a 75-mm mesh nylon membrane, the protoplasts were collected by centrifugation with 100 x *g* for 5 minutes at 4°C. These collected protoplasts were then subjected to two washes with a cold W5 solution (154 mM NaCl, 125 mM CaCl_2_, 5 mM KCl, 2 mM MES and 5 mM glucose). The protoplasts were rested on ice for 30 minutes, the protoplasts were centrifuged with 100 x *g* for 5 minutes at 4°C. and then resuspended in MMG solution (0.4 M mannitol, 15 mM MgCl_2_ and 4 mM MES). A mixture of 200 μL of the MMG solution containing approximately 10^x6^ protoplasts, 10 μg of each plasmid, and equal volume of a PEG solution (comprising 40% PEG 4000, 0.2 M mannitol, and 0.1 M CaCl_2_) was incubated at room temperature for 10 minutes. Subsequently, 1 mL of W5 solution was added to halt the transfection process, followed by centrifugation at 100 x g for 2 minutes. The protoplasts were resuspended in 1 mL of W5 solution and incubated in darkness at room temperature overnight. After the transfection, the mesophyll protoplasts were gently resuspended for microscopic examination.

### MeRIP-seq and data analysis

MeRIP-seq was conducted using a combination of methods from previously study^36^ and a commercial kit (NEB, E1610S). Briefly, Three-week-old plants were used for mRNA purification. Approximately 5 μg of mRNA mixed with 10 fmol of methylated and unmethylated standards (NEB, E1610S) was used for fragmentation (NEB, E6150S). RNAs were then used for anti-m^6^A (NEB, E1610S) and anti-GFP (Sigma-Aldrich, G1546) immunoprecipitation, respectively. Immunoprecipitates were washed with series of buffer as protocol suggested, and then eluted by incubation with 200 μL Proteinase K solution (50 mM Tris-HCl pH 7.5, 50 mM NaCl, 2 mg/mL Proteinase K, 1% SDS, and 10 U SUPERase-In RNase Inhibitor) on a Thermomixer shaker at 37°C for 1 h. RNA purified with Trizol / RNA Clean & Concentrator Kits (ZYMO, R1017) was then used for end-repair and 5' end ^32^P-γ-ATP labeling. Labelled RNA was purified with denature urea-gel and then conducted both two ends adaptor ligation. cDNA was synthesized with primer reverse complementary to the 3'end adaptor. PCR products amplified from RNA derived cDNA was subjected to sequencing on Illumina NovaSeq6000 platform in pair-end mode with 150 bp per reads at Novogene.

MeRIP-seq analysis was conducted according to a protocol ^57^. Briefly, raw data were initially filtered for a quality control and trimmed to remove the adaptor sequences by Cutadapt. Reads were aligned to *Arabidopsis thaliana* reference genome (TAIR 10) using HISAT2. To obtain high-confidence m^6^A peaks (HC-m^6^A) in each sample, exomePeak2 was conducted with threshold as P < 0.05 and log_2_(enrichment) > 1. Log_2_(enrichment) represents log_2_(IP/input), log_2_(fold change) represents log_2_(enrichment^mutant^/enrichment^wild-type^). To detect differential methylation levels, peak candidates calling by exomePeak2 with P < 0.99, log_2_(enrichment) > 0, and Deseq2 normalization were separated into m^6^A loci and background depending on whether or not the candidates overlapped with HC-m^6^A peaks, respectively. To discover methylation modification on pri-miRNAs, peaks were identified using exomePeak2 with pc_count_cutoff = 10, bg_count_cutoff = 5, P < 0.05, and log_2_(enrichment) > 0, or using sliding window algorithm with block of 25 nucleotides, IP_mean_count ≥ 5, winscore > 0, and normalization with median of genes^36^. The metagene analysis was performed with Guitar package. The motif of m^6^A peaks was analyzed using HOMER Motif Analysis. Integrative Genomics Viewer was used to visualize the m^6^A peaks.

### MeRIP RT-qPCR

MeRIP and RNA purification was conducted as above described. To detect m^6^A on pri-miRNA, full length mRNAs were used for IP. Subsequently, cDNA was synthesized with Oligo(dT) and random hexamers. Enriched m^6^A signals from IPs were initially normalized to their input and then to that of Col-0 for relative enrichment calculation.

### *In vitro* methylation assay

The RNA substrate sequence was adopted from a previous study^58^. The reaction mixture was performed in a 20 mM Tris-HCl pH 8.0, 100 mM NaCl, 2 mM MgCl_2_, 50 μM ZnCl_2_, 1 U RNasIn, 2 mM DTT, 6% glycerol, 0.1% NP-40 buffer with 2 μM RNA, 0.1 μM methyltransferase (MTA and MTB), 1 μM auxiliary protein (various forms of SE and His-sumo), 1 mM SAM (A4377, Sigma) in a total volume of 20 μL. Reaction was conducted at 37°C for 2 hours. Reaction was stopped by adding NaCl to 1 M and stayed at room temperature for 15 min. RNA was then purified using RNA Clean & Concentrator Kits (ZYMO, R1017) for subsequently MeRIP enrichment and ^32^P-γ-ATP labeling. Subsequently, RNA was visualized by running a 15% Urea-PAGE gel for phosphor imaging.

### Protein 3D structure stimulation

Protein structure prediction was performed using AlphaFold 2.2.0 with the alphafold2-monomer^29^ and alphafold2-multimer^30^ modules for protein monomers and complexes, respectively. The software package was downloaded from the official GitHub repository and the necessary dependencies were installed. The predicted structures were saved in PDB format in the output directory, and the model with the highest predicted score was used for subsequent analysis. The PDB files were visualized using ChimeraX.

### Graph drawing

Graphs with dot plots (individual data points) were drawn using GraphPad Prism 9. Models and illustrations were made with Biorender.

### Statistics and reproducibility

Statistical analyses of quantification results were performed using GraphPad Prism 9. All statistical data show the mean ± s.d. of at least three biologically independent experiments or samples. Two-tailed unpaired Student’s t-test, Fisher’s exact test, one-way ANOVA with Dunnett’s multiple comparison test, two-way ANOVA analysis with Dunnett’s multiple comparisons test, and two-way ANOVA analysis with Tukey’s multiple comparisons test were performed in the Graphpad Prism 9. DEseq2 was performed for identifying methylated peaks in MeRIP-seq analysis. The hypergeometric distribution test was conducted in R programming. No statistical method was used to pre-determine sample size, which was determined in accordance with previous studies^6,16,38^ and the standard practices in the field. All experiments were independently repeated more than three times with similar results obtained, and the exact replicate numbers are provided in the respective figure legends. No data were excluded from our studies. Plants were randomly assigned to experimental groups whenever possible. For other experiments, the experiments were randomized and the investigators were blinded to allocation.

## Reporting summary

Further information on research design is available in the Nature Research Reporting Summary linked to this paper.

## Data availability

All data are available in the main text or the supplementary materials. High-throughput sequencing data generated by this study can be accessed in the NCBI BioProject database under accession code <u>PRJNA1102430</u>. SE ChIP-seq data can be accessed in the European Nucleotide Archive (ENA) under accession number ERP016859^37^. The mta sRNA-seq data published in previous study can be accessed in the the NCBI GEO database under accession code GSE122528^23^. For pan-transcriptome analysis, data were downloaded from an Arabidopsis RNA-seq database, a comprehensive online repository housing over 20,000 publicly available Arabidopsis RNA-seq libraries (http://ipf.sustech.edu.cn/pub/athrna/)^26^. Genetic materials will be available when requested.

## Code availability

This paper does not report original code.

---

**Extended Data Figure 1.**
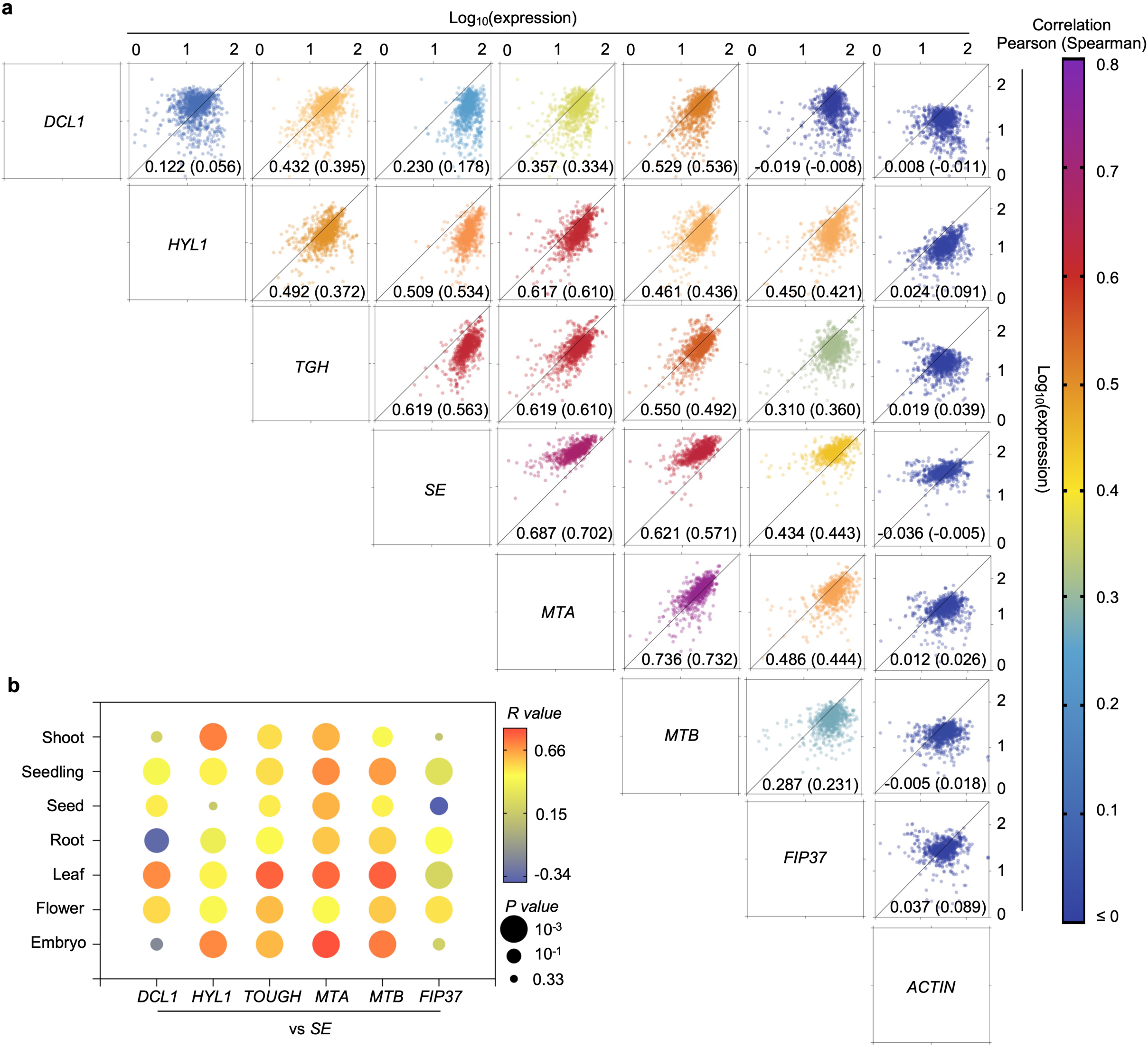

**Extended Data Figure 2.**
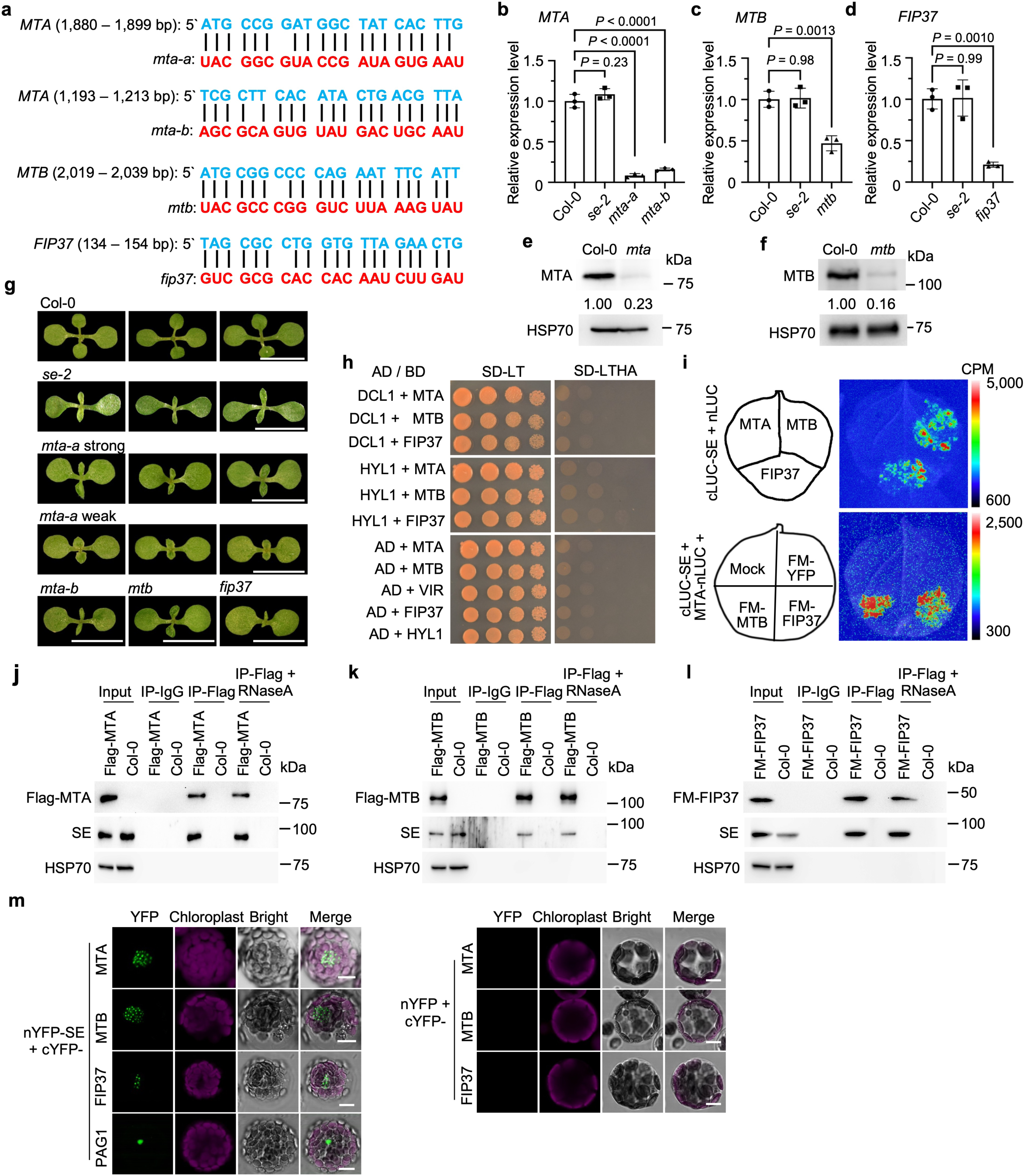

**Extended Data Figure 3.**
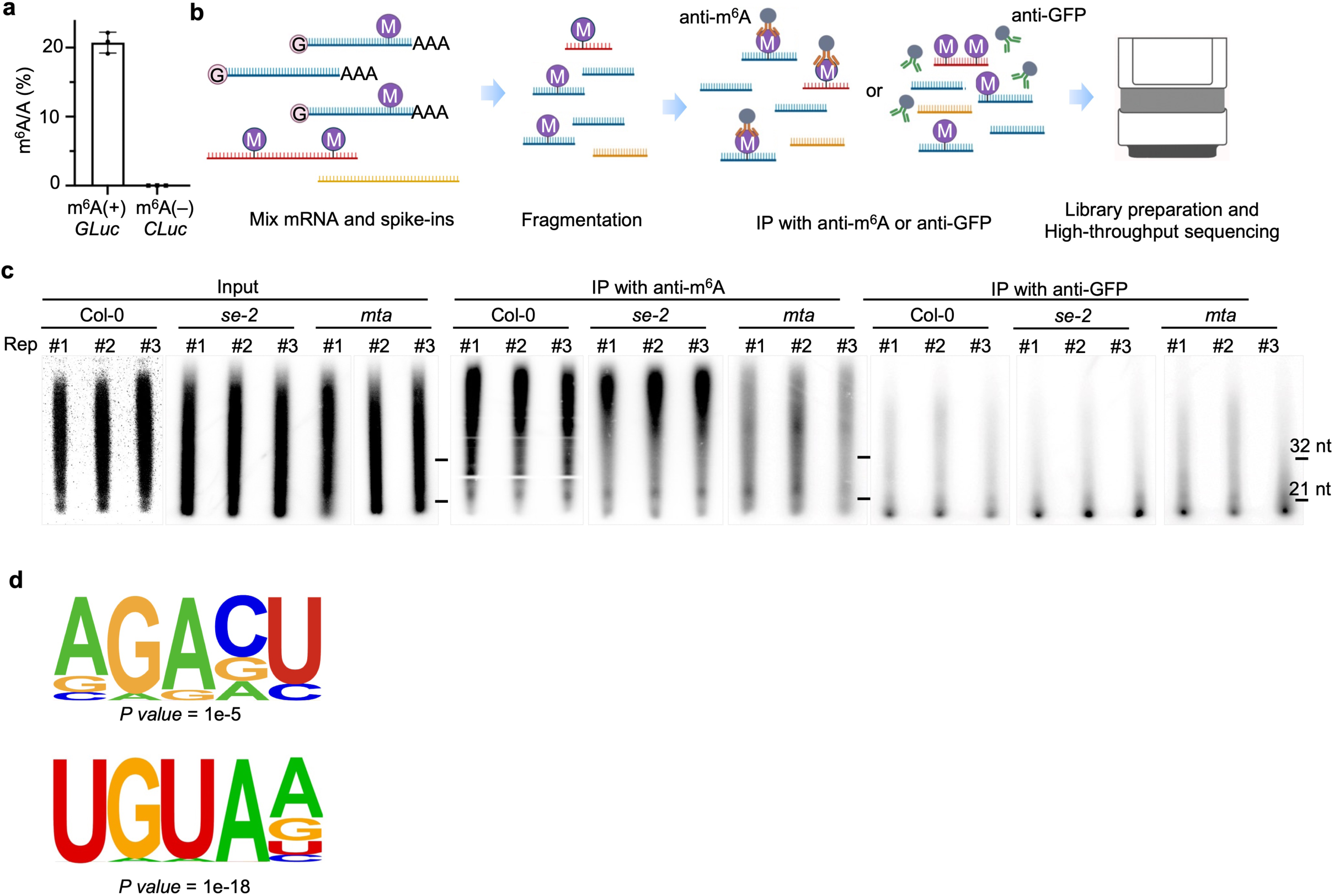

**Extended Data Figure 4.**
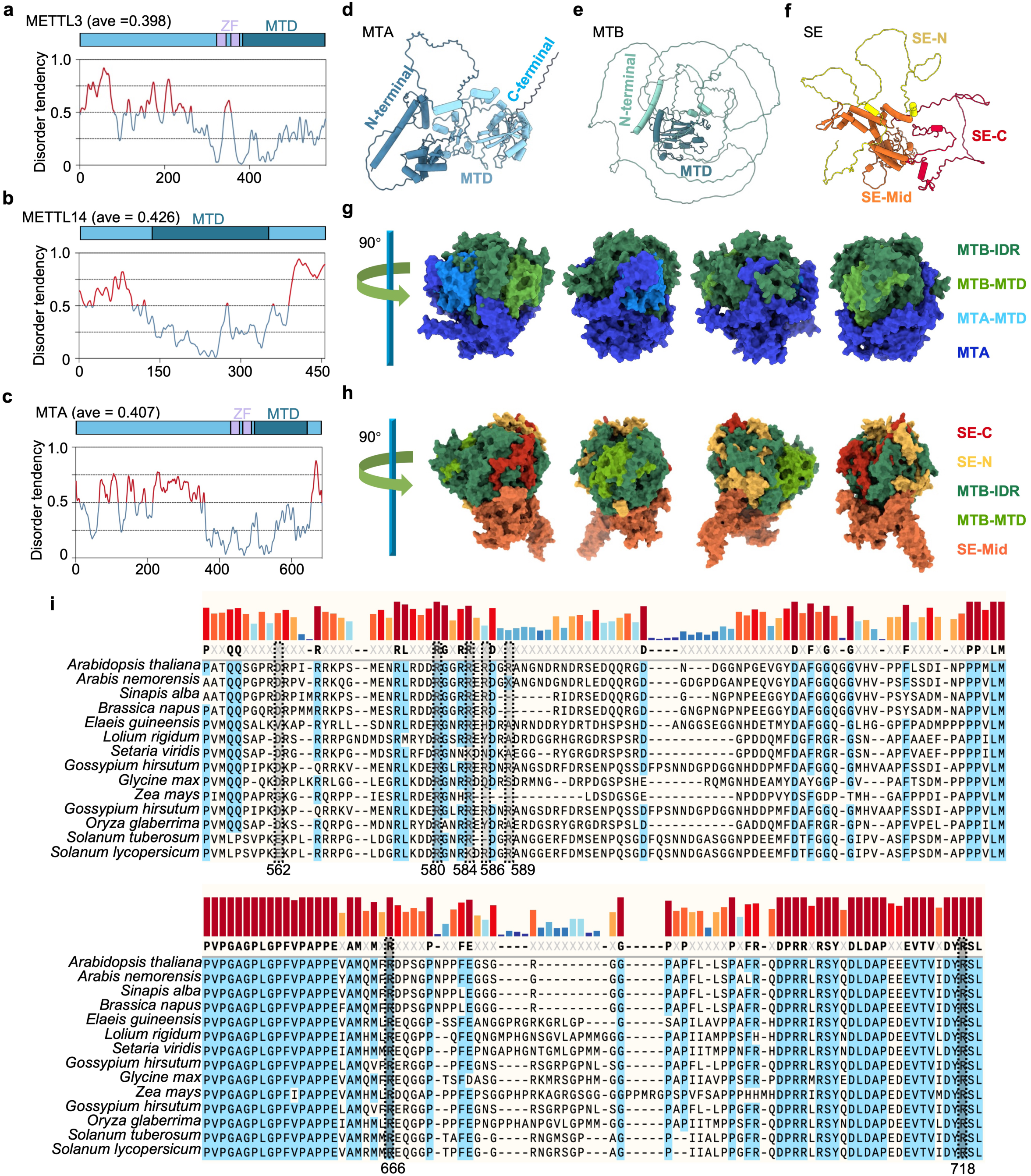

**Extended Data Figure 5.**
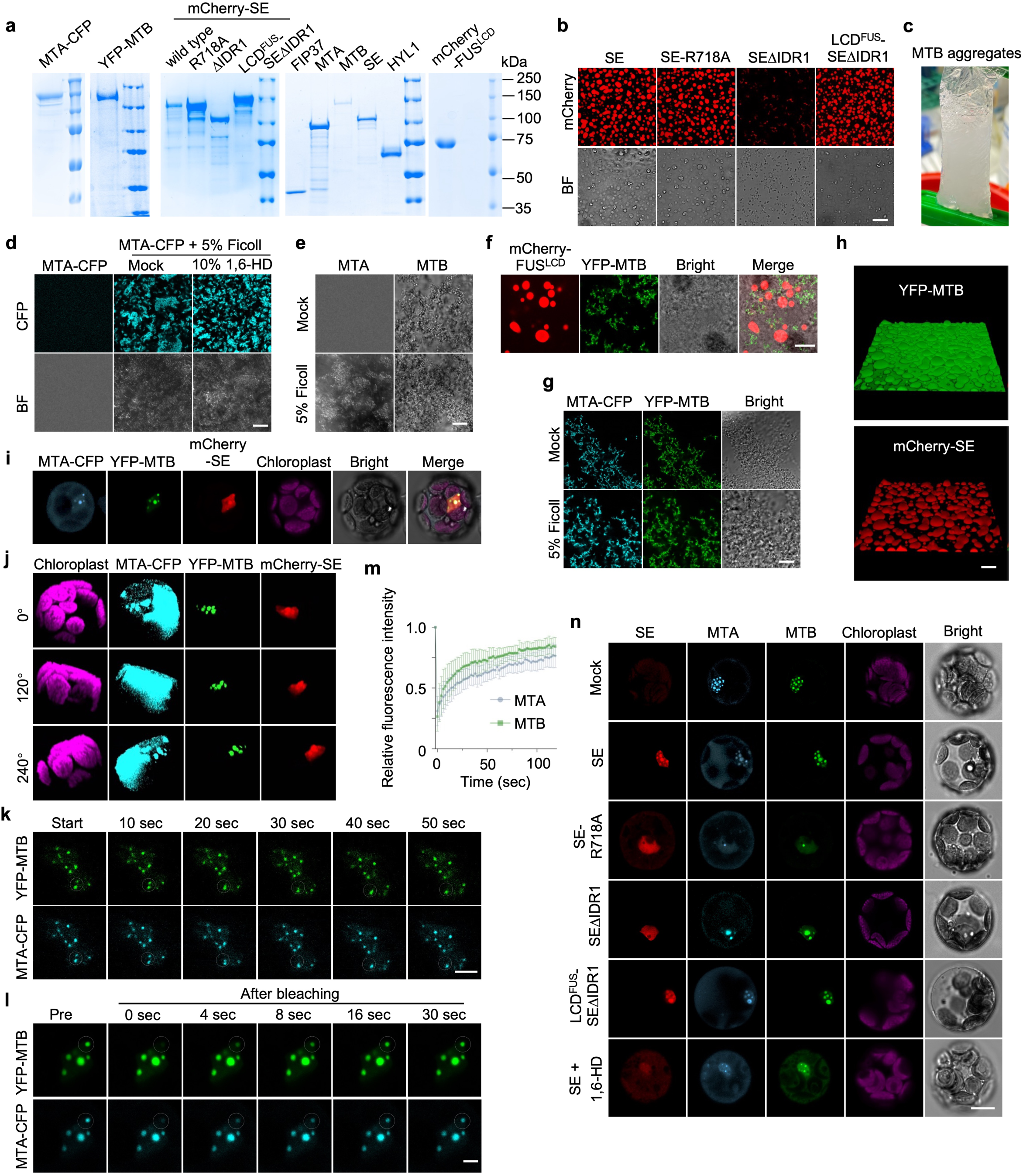

**Extended Data Figure 6.**
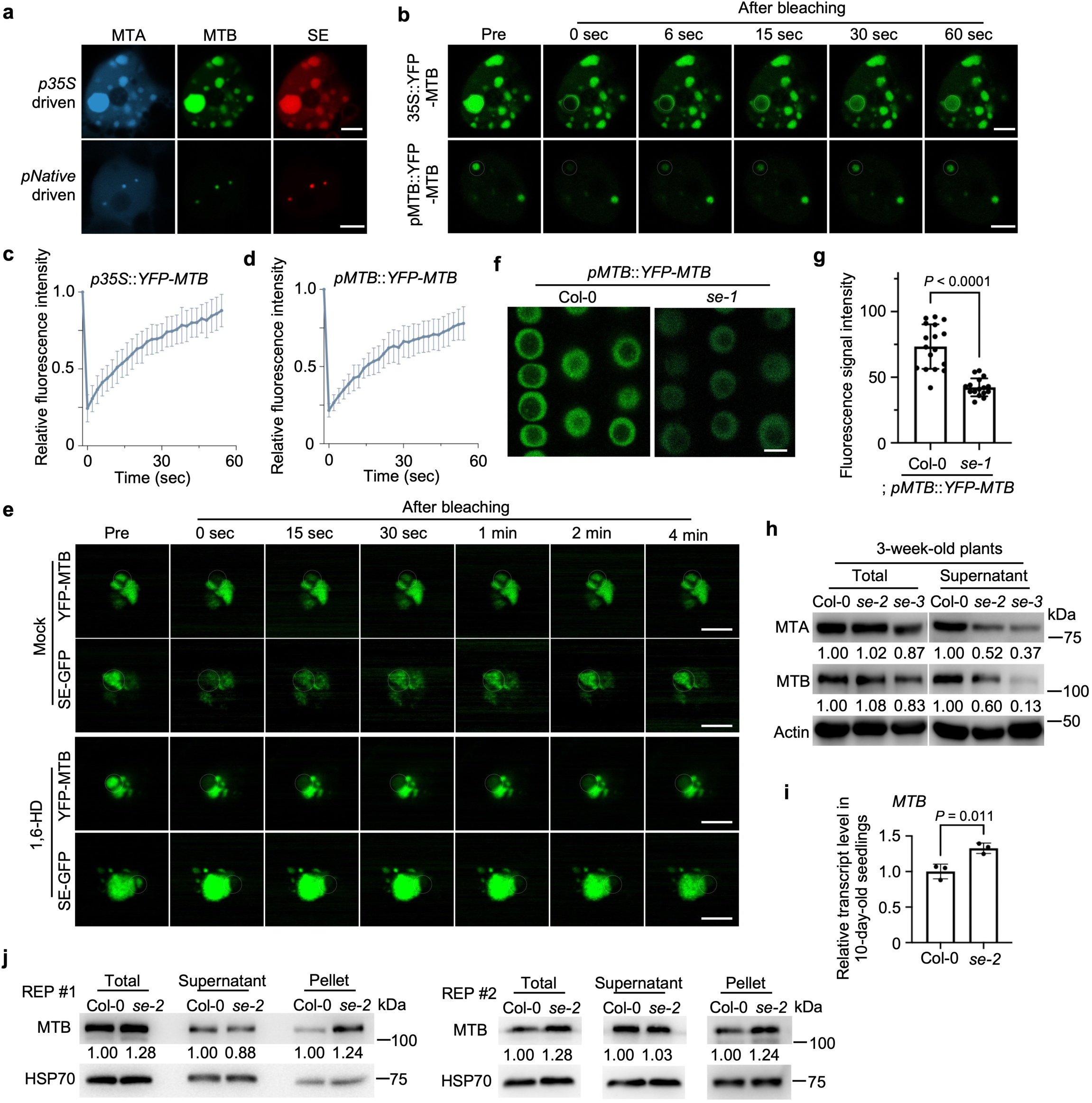

**Extended Data Figure 7.**
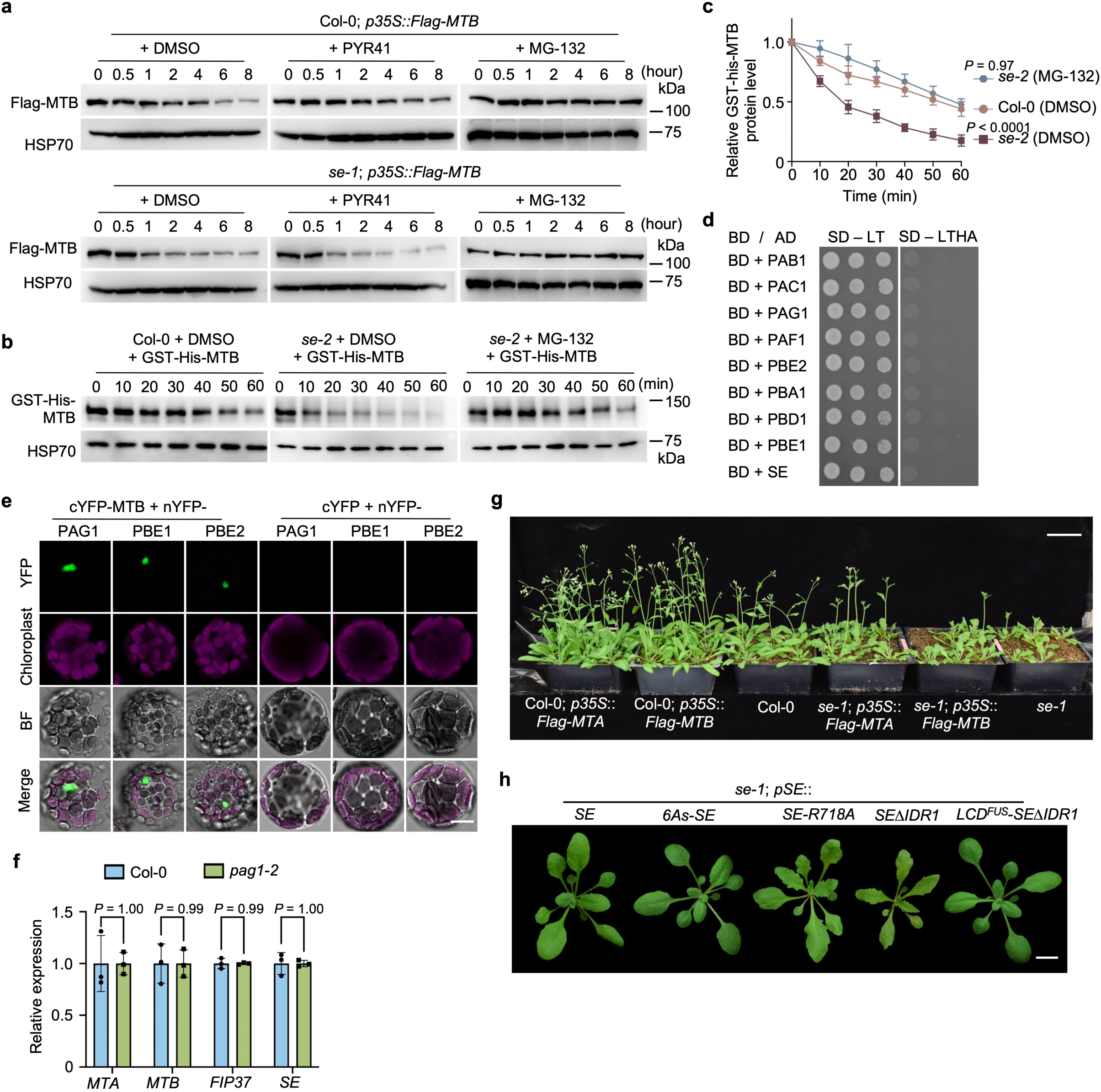

**Extended Data Figure 8.**
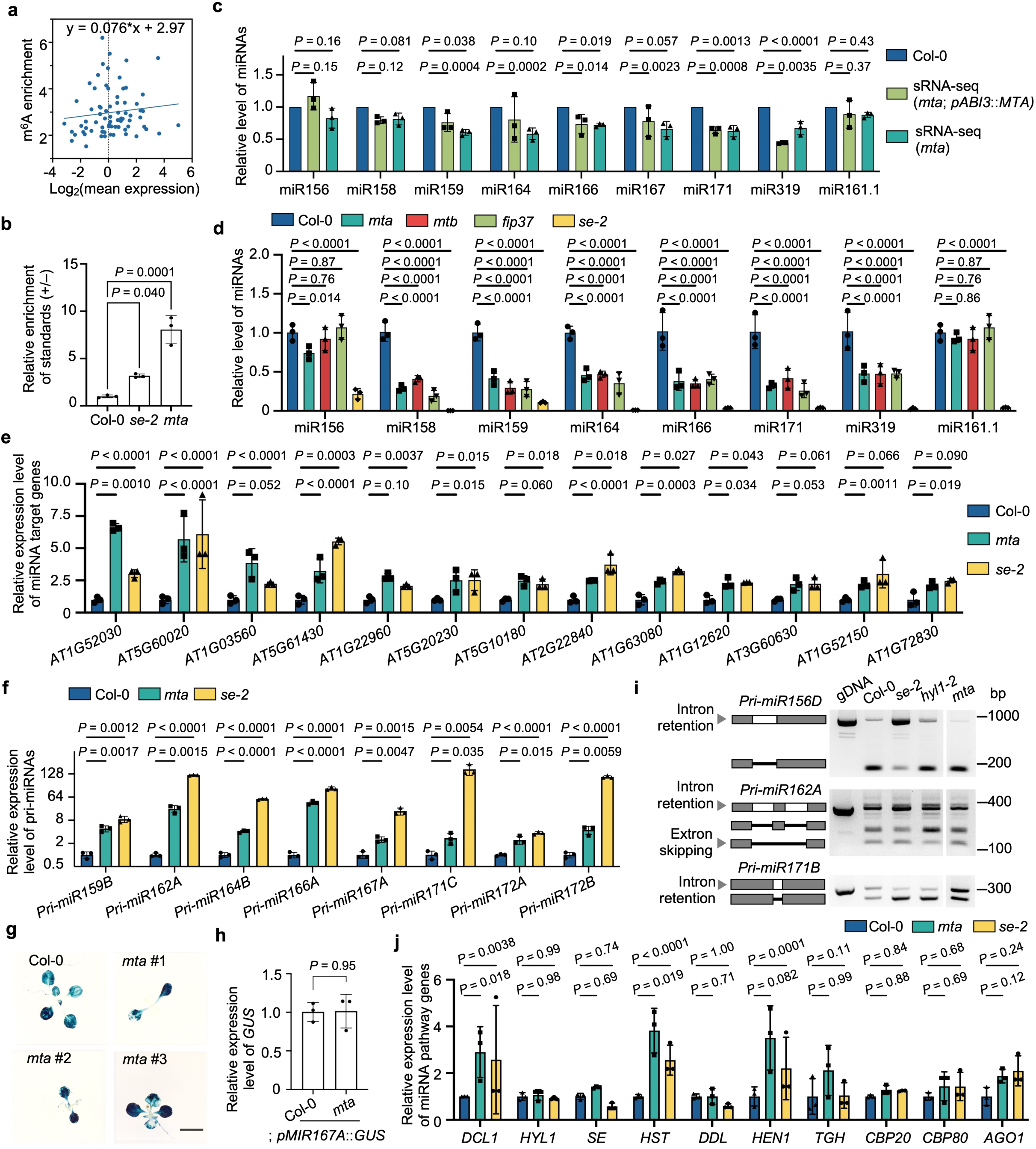

**Extended Data Figure 9.**
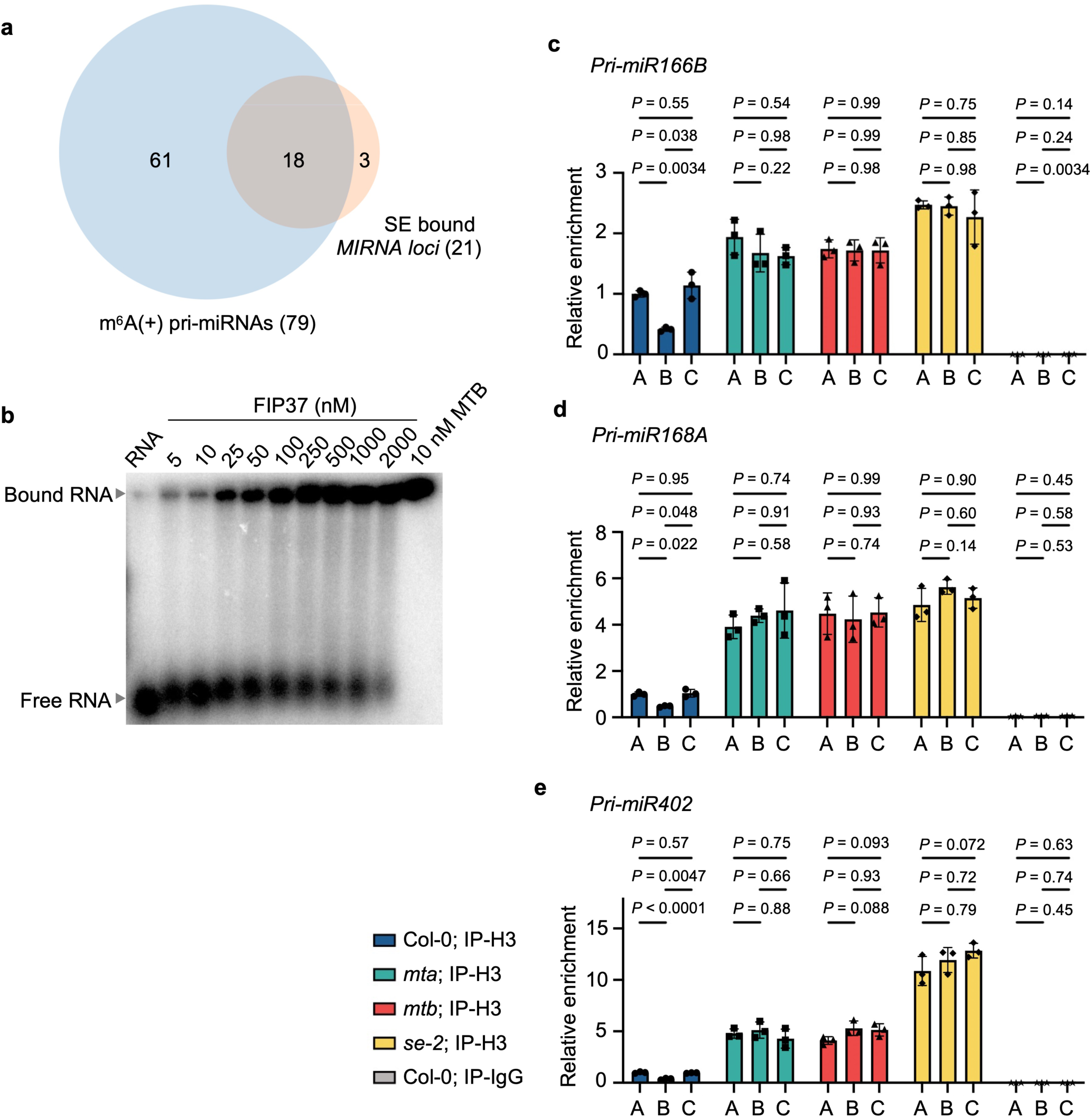

**Extended Data Figure 10.**
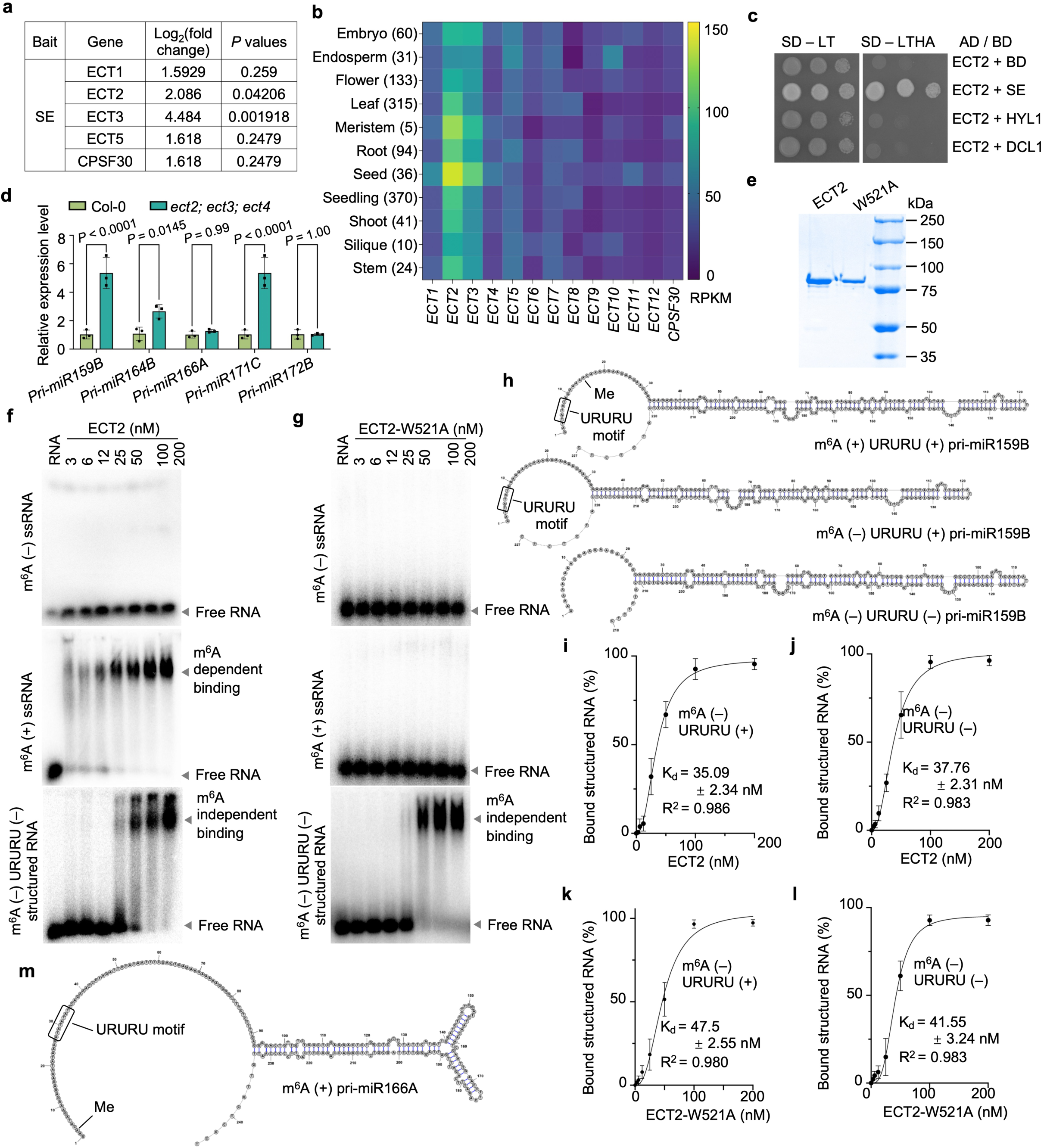

